# Navigating the peptide sequence space in search for peptide binders with BoPep

**DOI:** 10.1101/2025.01.20.633551

**Authors:** Erik Hartman, Firdaus Samsudin, Malcolm Siljehag Alencar, Di Tang, Peter J. Bond, Artur Schmidtchen, Johan Malmström

## Abstract

Peptides are short amino-acid chains that mediate essential biological processes, including antimicrobial defence, immune modulation and cell signalling. Their high degree of modularity, biocompatibility and capacity to bind proteins with high specificity make them attractive therapeutic candidates. However, identifying peptides that bind and modulate the function of specific proteins remains challenging due to the immense size of the peptide sequence space. To adress this challenge, we developed BoPep (**B**ayesian **O**ptimization for **Pep**tides), an end-to-end modular framework that effectively navigates the landscape of peptide-protein interactions by directing the search toward informative regions of sequence space and prioritizes candidates with high binding potential. By focusing computational effort where it is most informative and using calibrated uncertainty to balance exploration and exploitation, BoPep reduces the number of expensive docking evaluations by orders of magnitudes. We demonstrate the utility of BoPep by applying it to three sources of peptides: endogenous proteolytic fragments from clinical wound fluids, the complete human proteome, and a *de novo* design peptide landscape generated by diffusion-based backbone sampling. Using these sources, we uncover novel encrypted peptide classes that bind CD14 and identify peptides that neutralize the hemolytic activity of pneumolysin, a major bacterial virulence factor. Together, these findings show that BoPep accelerates the identification of testable therapeutic leads from large and diverse peptide collections. BoPep is available at GitHub.

## Introduction

Peptides are short chains of amino acids produced either through translation or through degradation of proteins. Their size and structure enable them to act as signalling molecules, antimicrobial agents and modulators of immune responses, roles that frequently depend on selective protein binding. These properties have made protein-binding peptides a focus area in biological research and a promising avenue for therapeutic development [1].

The number of possible amino acid configurations which constitute peptides is large, and the universe of possible sequences can be viewed as a landscape, where coordinates correspond to a unique arrangement of amino acids and reflect the associated properties and functions [2]. Recent studies have shown that organisms generate endogenous peptides through proteolytic processing of larger protein precursors [3–6], indicating that evolution has sampled parts of this peptide landscape. Some of these peptides exert immunological functions, supporting the notion that innate immune peptide effectors are embedded within diverse host proteins and can be released through proteolysis [7–21]. Despite the growing access of high quality peptidomic datasets, identifying functionally active peptides within the overwhelming background of inactive fragments remains a significant hurdle.

Recent advances have begun to make the amino acid sequence landscape more accessible for identifying peptide sequences with specific properties. Protein language models, for example, have learned the underlying grammar by capturing sequence patterns shaped by evolution [22]. Structure prediction models like AlphaFold, Boltz and ESMfold can infer three-dimensional structure and potential binding poses from amino acid sequences [23–26]. Generative models like RFdiffusion, BindCraft and BoltzGen sample the sequence space to generate structures with desirable characteristics [24,27,28]. Building on these advances, workflows such as EvoBind and RFpeptides have been developed specifically for peptide binder design [29,30]. Despite this progress, discovering functional peptides still resembles searching for a needle in a haystack. Thousands of candidates must typically be generated and screened through costly docking or heuristic scoring after generation, mirroring the inefficiencies of searching within the endogenous peptidome.

Here, we reasoned that deep learning models could aid in navigating the peptide landscape by gradually learning a relationship between binding and peptide representations. To implement this idea, we created BoPep (**B**ayesian **O**ptimization for **Pep**tides), an end-to-end modular framework that uses docking-derived features to train a probabilistic surrogate model and applies Bayesian optimization to balance exploration of uncertain regions with exploitation of high-performing candidates. This approach reduces the number of peptides that must be docked or co-folded by about tenfold and enables computationally efficient exploration of peptides within endogenous and *de novo* peptidomes. Here, we apply BoPep to identify CD14-binding peptides from two peptide sources, a previously characterized wound fluid peptidome and the complete theoretical peptidome encrypted in the human proteome. Further, we *de novo* designed and selected neutralizing peptides targeting pneumolysin (PLY), a significant virulence factor secreted by *Streptococcus pneumoniae* [31]. BoPep is available at GitHub under an MIT license.

## Results

Efficient discovery of peptide binders requires navigating a sequence space that is far too large to explore exhaustively and too costly to evaluate with docking. Bayesian optimization (BO) offers a principled approach for such settings. In BO, a probabilistic surrogate model predicts the outcome of an expensive evaluation, and this model is then used to identify which candidates are most valuable to assess. In this way, the search can be directed either toward candidates predicted to score highly or toward regions where uncertainty is high and additional information would be most informative. The objective is to identify promising candidates while evaluating only a small fraction, since the computational overhead of updating the surrogate model is minor compared with the cost of evaluation.

BoPep encompasses all components required to apply BO to peptide binder discovery, including modules for embedding, docking, scoring and surrogate modelling. Further, it provides methods that string these modules together to search a predefined peptide library, or complete proteomes. It also provides a workflow for *de novo* binder design in which large libraries of candidate peptides are generated before the optimization begins (**Fig. 1a**). The BO procedure starts by sampling peptides from a dataset and docking them to the pre-defined target. These initial evaluations are used to train a surrogate model that estimates binding potential and serves as a computationally efficient substitute for direct docking. Based on these predictions, a new set of peptides is selected for evaluation, and the cycle is repeated for a fixed number of iterations. By concentrating evaluations on informative regions of sequence space, BoPep reduces the computational burden of docking and co-folding (see **Methods: Computational efficiency**). In the following sections we describe the four core modules and how they are integrated within this framework.

**Fig. 1.**
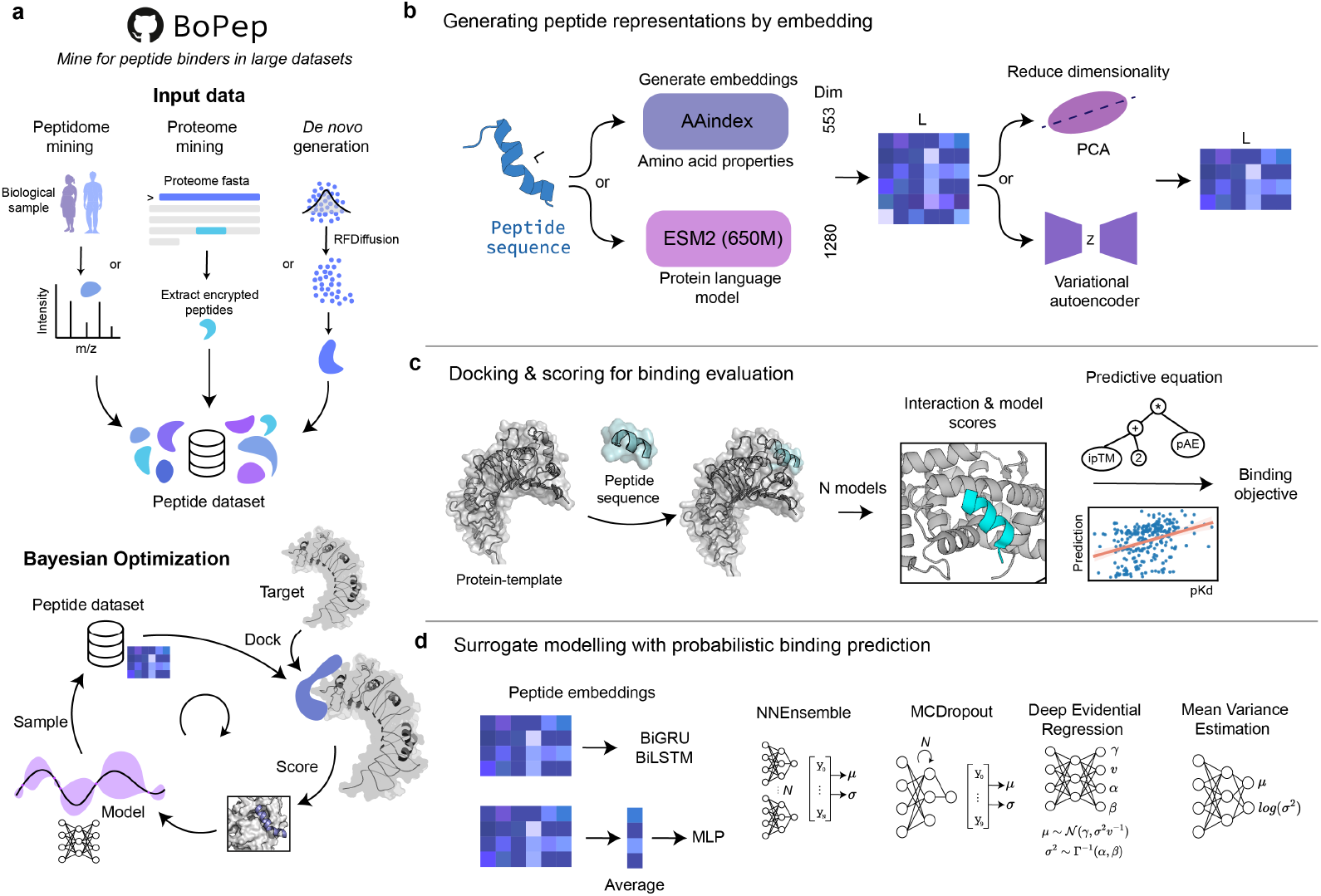
Overview of the BoPep framework for peptide binder discovery. **a** BoPep starts with a set of peptide candidates, which can be derived from any source, such as through endogenous peptide mining of biological samples, complete proteomes, or from a *de novo* binder generation pipeline. The process begins by embedding the peptides into a latent space, which is followed by an iterative process, involving docking peptides against a target, scoring the binding interactions, and training a surrogate model. The surrogate model is then used to guide further sampling by selecting which peptides should be selected in the next iteration. **b** Peptide sequences are embedded into numerical representations using amino acid properties (AAindex, 553 dimensions) or protein language models (ESM2, 650M parameters, 1280 dimensions). To reduce their dimensionality, PCA or a VAE can be employed, yielding lower dimensional embeddings. **c** Docking and scoring of the peptides to the target protein constitutes the computationally expensive evaluation function. After structural modelling of the peptide-protein complex, interaction metrics and confidence scores are combined in a predictive equation to approximate binding affinities (pKd). The predictive equation is shown in Fig. 2. **d** Surrogate modelling is used to predict binding scores and guide the search. BoPep includes two types of architectures of four modalities for uncertainty estimation. The architectures include recurrent architectures (BiGRU, BiLSTM) or multilayer perceptrons (MLPs), and probabilistic approaches including neural network ensembles, Monte Carlo dropout, deep evidential regression, or mean-variance estimation. These models provide binding predictions with associated uncertainty estimates, enabling BO.

A useful BO procedure requires a representation of peptide sequences that captures functional similarity between peptides and allows the search to move through sequence space in a meaningful way. BoPep therefore embeds peptides either through physicochemical descriptors from AAindex [32] or through contextual sequence representations learned by the protein language model ESM2 [22]. These high-dimensional embeddings can be reduced with principal component analysis (PCA) or variational autoencoders (VAEs) to obtain continuous latent spaces that are easier to explore during optimization (**Fig. 1b**).

To evaluate candidate binders, BoPep performs structural modelling of peptide–protein complexes using co-folding, also referred to as docking, with AlphaFold2, implemented in ColabFold [33], or with Boltz-2 [26] (**Fig. 1c**). The resulting structures are assessed with confidence-based and energy-based metrics, and these scores provide the training data for the surrogate model. Details of the scoring procedure and the construction of a binding prediction equation are provided in **Results: A predictive equation for peptide binding prediction**.

The surrogate model predicts the binding scores from the embeddings and quantifies uncertainty, enabling a strategic traversal of the sequence space. BoPep supports multilayer perceptrons and recurrent architectures, along with several methods for estimating predictive uncertainty (**Fig. 1d**). These include Monte Carlo (MC) dropout, model ensembles, mean-variance estimation (MVE) and deep evidential regression (DER), which together provide the estimates of confidence needed to guide exploration and exploitation during optimization [34,35].

Together, these modules form a modular and extensible framework that links peptide representation, structural evaluation and uncertainty-aware modelling into a coherent optimization loop. By reducing dependence on exhaustive docking and focusing evaluations on informative regions of sequence space, BoPep directly addresses the challenge of discovering peptide binders within a vast and sparsely explored landscape.

### Identification of a predictive equation for peptide binding

During optimization we require an *objective* that reflects how likely a peptide is as a potential binder. Direct prediction of binding probability and affinity remains difficult and is one of the central challenges of structural biology. Recent models trying to tackle this problem are impractical and not feasible for implementation in BoPep. However, there are several proxy measures that are available through the structural prediction models or that can be cheaply computed from a single predicted complex, using e.g. Rosetta-based tools [36]. These metrics capture co-folding confidence, geometry and physics-based scores, which are typically used to filter candidates in binder generation pipelines. On their own they are not reliable predictors of binding, however, when combined these represent a suitable quantity for guiding optimization tailored to peptide-protein complexes. We therefore set out to use symbolic regression [37] to generate a predictive equation that could be used as a proxy for binding probability and affinity.

To generate a predictive equation we first assembled a dataset of peptide-protein complexes and accompanying pKd values. We assembled such a dataset by filtering complexes from PDBbind resulting in 300 linear peptide-protein complexes [38] (**Fig. 2a**). For each template structure, we generated an equivalent set of structural decoys, either by residue shuffling or by generating random peptides, yielding balanced true and decoy sets. All complexes were re-predicted with AlphaFold2 multimer and scored with a series of simple metrics available in BoPep, resulting in a feature matrix that captured energetic, confidence based and structural descriptors.

**Fig. 2.**
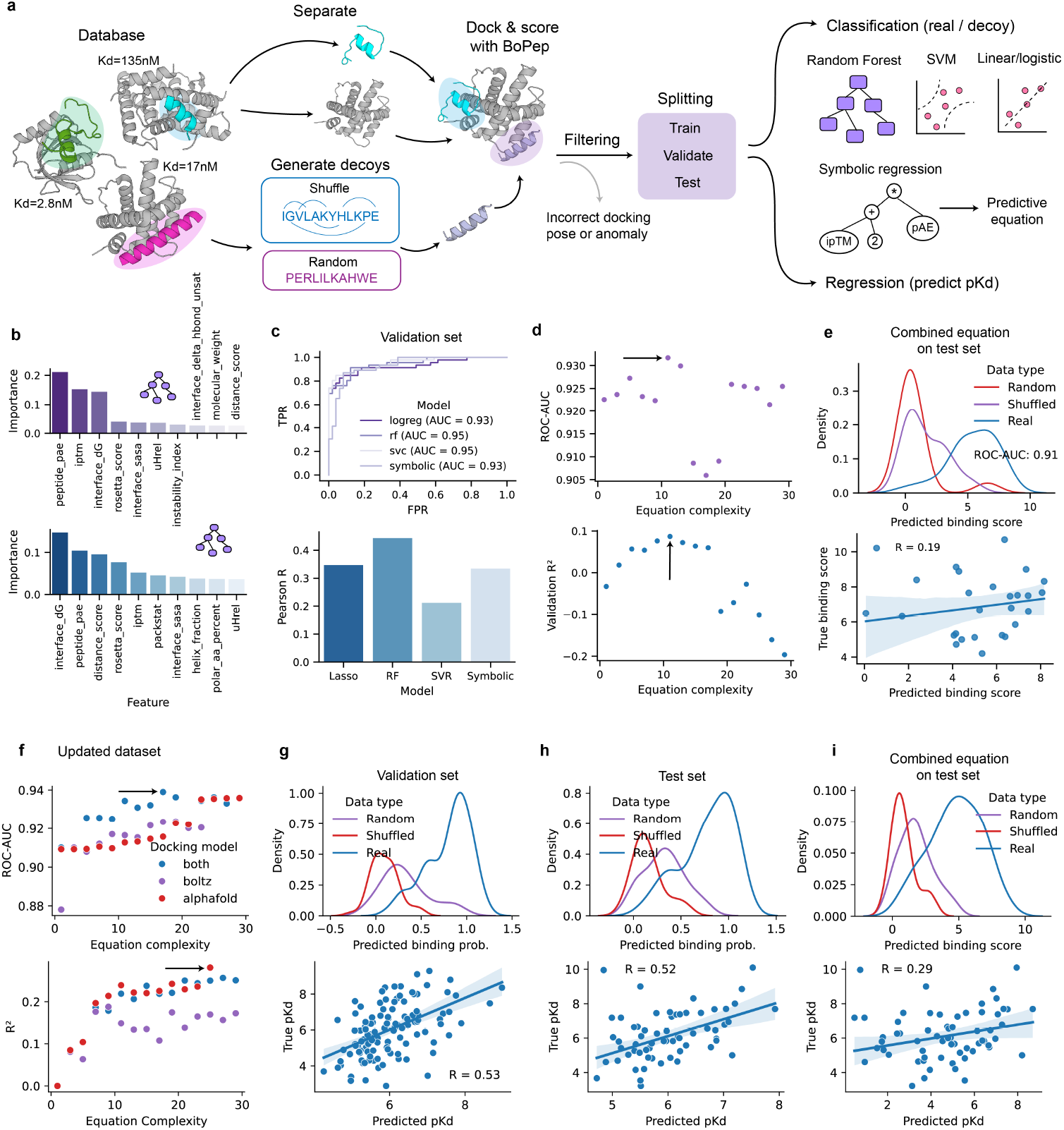
Generating a predictive proxy objective for binding affinity and probability. **a** To generate a binding score that can be used as the objective during optimization, linear peptide-protein complexes with measured affinities were extracted from PDBbind. For each complex a decoy was created by residue shuffling or by generating random peptide sequences. All peptides were docked onto the target template after removing the peptide chain, and scored with BoPep. The complexes were then filtered and split into training, validation, and test sets. Downstream analyses involved classification of real versus decoy binders (random forest (RF), support vector classifier (SVC), symbolic regression) and regression to predict affinities (pKd). **b** Feature importances of the ten most important features of random forest models for classification (top) and regression (bottom). **c** Model performance on the validation set. Top: ROC curves comparing logistic regression, random forest, SVM and symbolic regression classifiers. Bottom: Pearson correlation between predicted and experimental pKd for regression models (lasso, RF, support vector regressor (SVR), symbolic). **d** Several equations were generated using symbolic regression and the panels show the effect of equation complexity on predictive performance. Each point represents an equation. Classification accuracy (ROC-AUC, top) on the validation set was almost constant across model complexity while regression performance (*R*^2^, bottom) peaked at around complexity 10. The arrows point to the chosen equations which were evaluated both based on performance. **e** Final evaluation of combined expressions on the test set. The top panel shows the distributions of combined scores for real and decoy binders and the bottom shows a linear regression of the predicted and actual pKd. Panels **f-i** show results for the updated dataset. **f** Symbolic expression complexity and accuracy on an updated dataset. The arrows point to the chosen equations. **g** The performance of the updated equations of regression and validation on the validation set. **h** The performance of the updated equations of regression and validation on the test set. **i** The performance of the combined equations of regression and validation on the test set.

The importance of the features were estimated with random forests, demonstrating that a mix of confidence scores, biophysical scores and shape complementarity were predictive of binding probability and affinity (**Fig 2b**). These features are commonly used when filtering candidate binders in generative design workflows and have been shown to correlate with binding affinity in earlier studies [24,27,30]. Using PySR, we generated several equations of varying complexity and selected the optimal expressions based on validation loss (**Fig. 2d**). Their performance was similar to that of other machine learning models despite their simplicity and interpretability (**Fig. 2c**).

The classification equation involved only the ipTM and the peptide predicted aligned error (pAE),

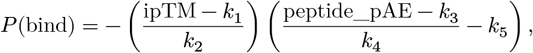

whereas the regression equation incorporated the Rosetta score, the interface ΔG and a distance based packing metric,

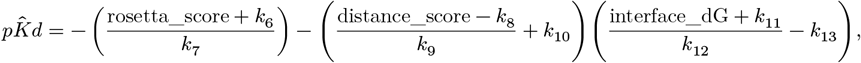

where *k*_*i*_ are constants (see Methods for constant values).

We combined these expressions by multiplying 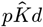 and *P* (bind) to form a composite predictive score that prioritized candidate binders in the optimization experiments described below (**Fig. 2e**). The score can be updated easily and users may substitute their own metrics or objective functions. During this project the PDBbind database expanded, increasing the number of available linear peptide-protein complexes. Repeating the symbolic regression with the enlarged dataset produced an expression with improved predictive power and revealed that AlphaFold derived features were more informative than those from Boltz 2 (**Fig. 2f-i**). All equations are available in the BoPep repository.

### Benchmarking the optimization hyperparameters

Employing BoPep involves many design choices, including how peptides are embedded, how the surrogate model is constructed and how predictive uncertainty is estimated, among other parameters. We therefore performed systematic benchmarks of embeddings, surrogate architectures and uncertainty modalities using a controlled subset of peptides drawn from clinical wound fluid peptidomes (see **Fig. 4** for dataset details).

We first assessed peptide embeddings derived from amino acid property indices (AAindex) and from a protein language model (ESM2) (**Fig. 3a**). To compare their intrinsic information content, we quantified how many principal components were required to retain 95 percent of the variance in each embedding type. ESM embeddings required more components than AAindex in both sequence-averaged (1D) and position-resolved (2D) settings (**Fig. 3b**), consistent with the higher representational complexity of the language model. Visualization of raw and reduced embeddings using UMAP showed that PCA largely preserves the global structure of the embedding space, whereas the latent space in the variational autoencoder (VAE) is re-organized (**Fig. 3c**).

**Fig. 3.**
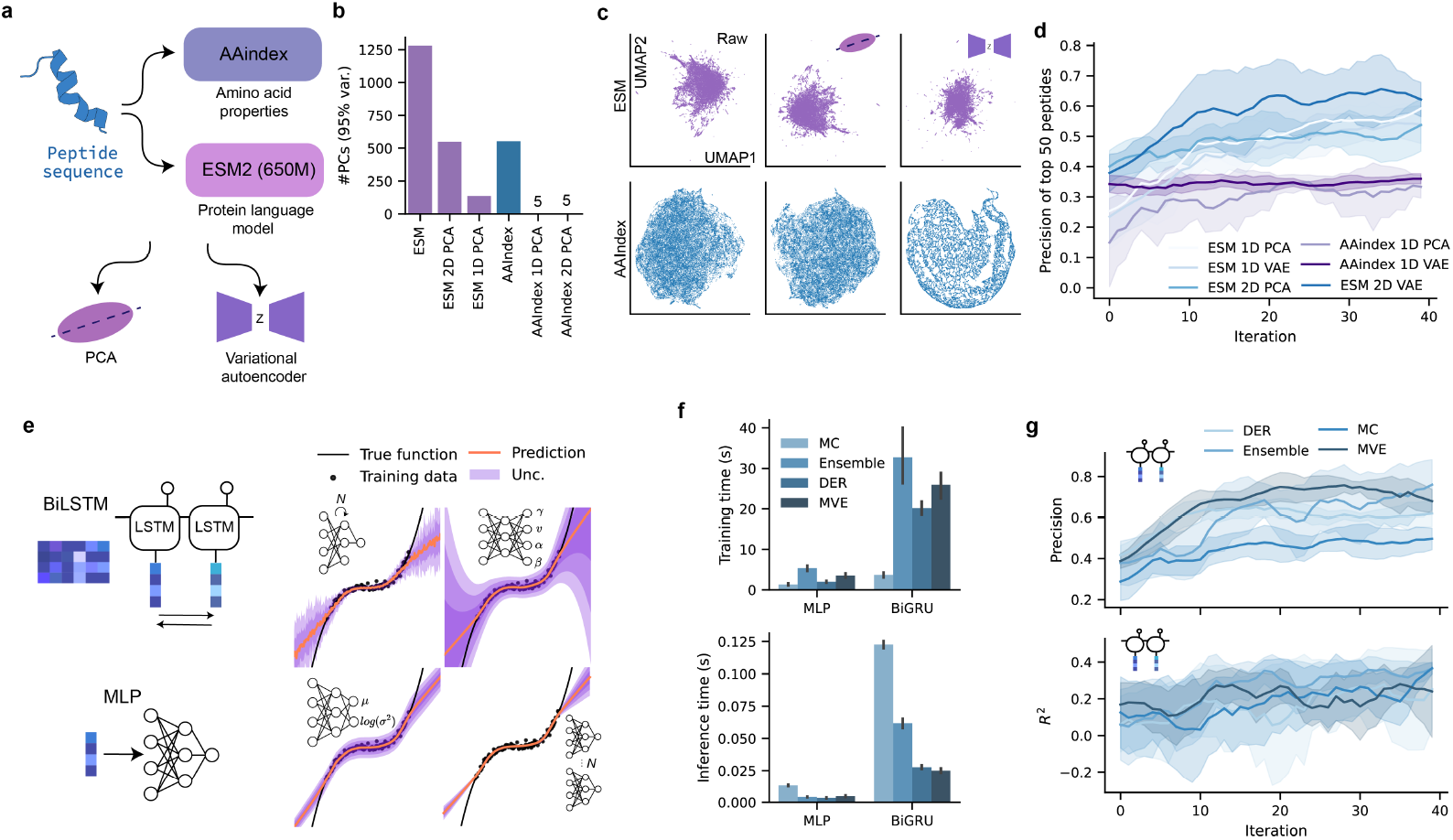
Benchmarking peptide embeddings, surrogate architectures and uncertainty estimation in BoPep. **a** Peptide sequences are embedded either using amino acid property indices (AAindex) or a protein language model (ESM2, 650M parameters). The resulting high-dimensional representations are reduced using principal component analysis (PCA) or a variational autoencoder (VAE) to obtain low-dimensional latent embeddings used by the surrogate model. **b** Number of principal components required to retain 95% of the variance for different embedding configurations. **c** UMAP projections of raw and reduced embeddings. PCA largely preserves the global structure of the embedding space, whereas VAE compression yields an approximately Gaussian latent space. Top row, ESM embeddings; bottom row, AAindex embeddings. **d** Optimization performance during a limited BoPep search using a fixed test bed of 500 docking results generated by the BoPep initializer. The precision of identifying the top 50 peptides is shown across optimization iterations for different embedding and reduction strategies. The lines are smoothed using a rolling window of size 10. Shaded regions indicate ±1 s.d. across triplicate runs. **e** Illustrations of surrogate model architectures and probabilistic uncertainty modalities. Multilayer perceptrons (MLPs) operating on sequence-averaged (1D) embeddings are compared to recurrent architectures (BiLSTM/BiGRU) applied to position-resolved (2D) embeddings. Predictive uncertainty is estimated using Monte Carlo (MC) dropout, deep evidential regression (DER), model ensembles and mean–variance estimation (MVE). The approaches are illustrated on a synthetic regression task with inputs sampled as *x* ∼ [−4, 4] and targets defined as *y* = *x*^3^ + *ϵ*, where *ϵ* ∼ 𝒩(0, 3). **f** Training time (top) and inference time (bottom) are shown for MLP and BiGRU architectures combined with different uncertainty modalities. Bars represent mean values and error bars indicate ±1 s.d. **g** To compare uncertainty modalities, benchmarking optimizations were performed with the different modalities when applied to the BiGRU architecture on the ESM embeddings. Top, precision of identifying the top-ranking peptides; bottom, *R*^2^ of surrogate predictions across optimization iterations. The lines are smoothed using a rolling window of size 10. Shaded regions indicate ±1 s.d. across triplicate runs.

To assess how factors influence optimization behaviour, we monitored various performance metrics during limited BoPep searches applied to 500 docking results generated using the BoPep initializer. In these benchmarks, the optimization objective was to find the peptide that maximizes ipTM, and metrics like the recall and precision of the top 50 peptides were monitored.

In the first comparison, we evaluated the effect of embedding strategies. 1D AAindex embeddings underperformed, whereas the remaining embedding configurations showed broadly similar behaviour across iterations (**Fig. 3d**). The poor performance of 1D AAindex is consistent with its construction from fixed per-residue property indices that do not adapt to sequence context, limiting discriminative power when embeddings are averaged across positions.

We next benchmarked surrogate model architectures and uncertainty modalities (**Fig. 3e**). Recurrent architectures required more time overall than MLPs (**Fig. 3f**). The benchmarks were run in triplicates and concluded that ESM embeddings in combination with BiGRU architectures and DER or MVE uncertainty modalities were optimal (**Fig. 3g**). Due to the ability of DER to estimate aleatoric and epistemic uncertainty, we continued with DER for future experiments.

Taken together, BoPep integrates a predictive equation for ranking binders with state-of-the-art embedding strategies and probabilistic surrogate modelling. We applied this framework to diverse peptide collections to evaluate the ability to recover bioactive peptide binders from large endogenous and *de novo* design peptide landscapes.

### Mining clinical proteomes for encrypted immunomodulatory peptides binding to CD14

To evaluate the performance of BoPep in a biologically relevant setting, we applied it to search for encrypted immunomodulatory peptides that bind to CD14. The N-terminal domain of CD14 mediates transfer of lipopolysaccharide to MD-2 and activation of the TLR4 pathway, a central driver of inflammatory responses [11,39] (**Fig. 4a**). Blocking this interaction attenuates excessive inflammation making this interaction a promising immunomodulatory target. Here, we used a previously characterized wound fluid peptidome as a starting point to assess whether BoPep can identify candidate binders to CD14 from a large and heterogeneous experimentally obtained endogenous peptide pool.

**Fig. 4.**
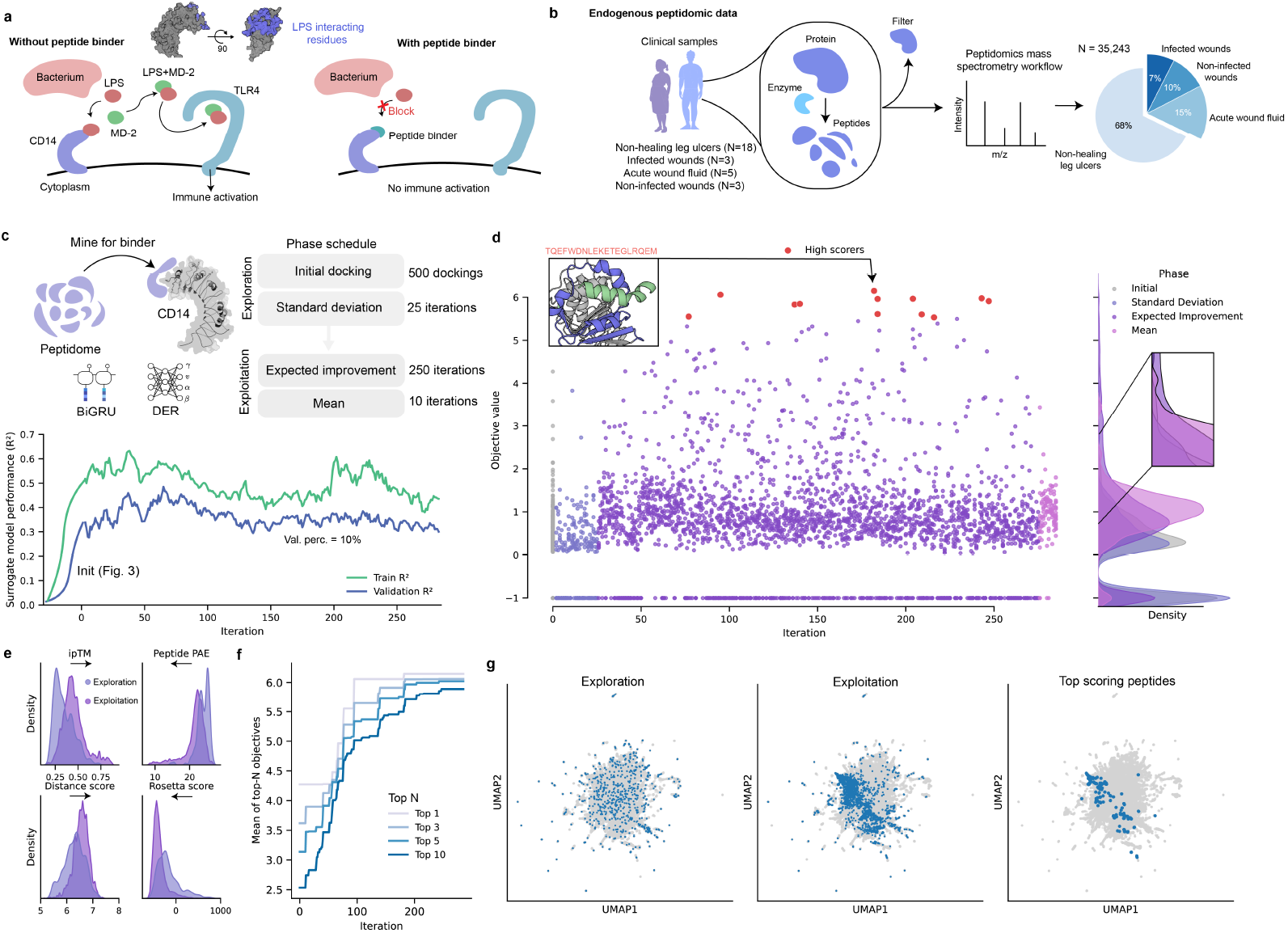
Identifying endogenous peptide binders to CD14. **a** Lipopolysaccharide (LPS) activates the innate immune receptor TLR4 through interactions with MD-2 and CD14. A schematic illustration how peptide binder blocks CD14-LPS interactions preventing downstream immune activation. **b** Endogenous peptide candidates were obtained from clinical peptidomic datasets comprising non-healing leg ulcers (n = 18), infected wounds (n = 3), acute wound fluid (n = 5), and non-infected wounds (n = 3), yielding 35,243 unique peptides. **c** The search was conducted using a BiGRU DER model, and embeddings were generated with ESM followed by PCA for dimensionality reduction. An initial docking round using 500 peptides seeded the surrogate model. The schedule started with exploration (sampling peptides with high uncertainty) followed by exploitation (selecting peptides with high predicted binding scores). Model performance (*R*^2^) is shown for training (80% of data) and validation sets (20% of data). The curves were smoothed computing running averages with a window size of 5. **d** Each dot represents a peptide, colored by the acquisition function used in each phase. The peptides with anobjective value > 5 are colored red for clarity. Right: density distribution of acquisition scores. The inset highlights the region where results from the exploitation demonstrating a clear shift compared to the results from exploration. **e** Distributions of docking and scoring features (ipTM, peptide pAE, distance score, Rosetta interface energy) across candidate peptides. **f** The mean objective value of the top-ranked peptides across iterations. **g** UMAP projections of the sequence space. Exploration sampled broadly across the embedding space (left), exploitation concentrated on predicted binders (middle), and top-scoring peptides localized to a compact region (right).

The clinical peptidomic datasets comprising non-healing leg ulcers (n = 18) [4], infected wounds (n = 3), acute wound fluid (n = 5), and non-infected wounds (n = 3) [14] yielded 35,243 unique peptide sequences (**Fig. 4b**). An initial docking round of 500 peptides seeded the BoPep surrogate model (same results as in the benchmark, *Fig. 3h*), which then guided iterative exploration and exploitation of the remaining space (**Fig. 4c**). In total 9.5% of the search space was evaluated with docking. Model performance stabilized after approximately 150 iterations, with validation *R*^2^ values maintained around 0.3-0.4 (**Fig. 4c**).

The optimization trajectory revealed a progressive enrichment of high-scoring peptides (**Fig. 4d**). Feature distributions of docking score components (ipTM, peptide pAE, distance score, Rosetta interface energy score) highlight how exploitation results in better binder candidates (**Fig. 4e**). The objectives of top binders increased until plateauing at 200 iterations, indicating that the search converged after docking ∼7% of the dataset (**Fig. 4f**). UMAP projections of the exploration and exploitation phase reveals that exploration covered the peptide space broadly, while exploitation concentrated predictions in tighter regions where the top-scoring peptides were located (**Fig. 4g**). These results demonstrate that BoPep can identify peptide–protein binders to central inflammatory mediators in a flexible and targeted manner using endogenous peptide repertoires. In the case of CD14, the framework isolated candidate binders that provide a starting point for the rational design of peptides capable of modulating TLR4 activity.

### Helical structure underlies the preferential binding of endogenous peptides to CD14

Inspection of the peptides with an objective value greater than four from the BoPep search for CD14 binders revealed an enrichment of peptides derived from proteins with high *α*-helical content, including apolipoproteins and several cytoskeletal proteins (**Fig. 5a,b**). These findings prompted us to investigate whether AlphaFold was biased towards peptides encrypted in proteins with high structural confidence. Further analysis confirmed that protein helicity, peptide region helicity and backbone confidence (pLDDT) were correlated with docking scores (ipTM and interface ΔG). Furthermore, cumulative recovery curves showed that the highest-scoring binders arose almost exclusively from helical, high-confidence regions (**Fig. 5c**).

**Fig. 5.**
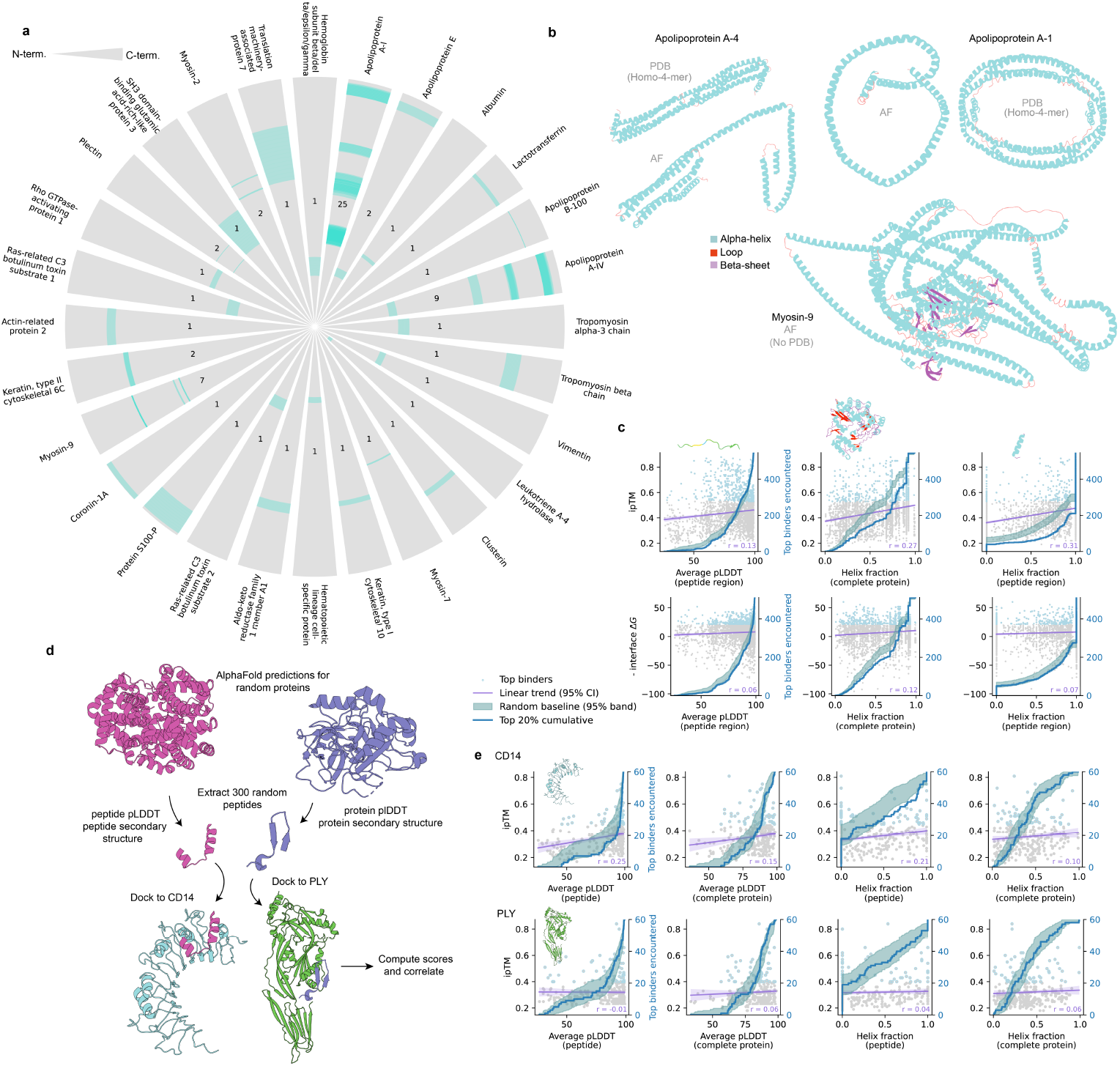
Basis of helix enrichment among CD14 peptide binders. **a** Circular plot of the source proteins for the top-ranked CD14 candidate binders identified by BoPep. Many peptides (regions highlighted in cyan) originate from proteins with high *α*-helical content, such as apolipoproteins, myosins, tropomyosins and vimentin. The numbers indicate peptide counts per protein and the highlighted regions show the protein regions where the peptides originate from in the protein backbones. **b** Representative structures of source proteins predicted by AlphaFold (AF) or obtained from the Protein Data Bank (PDB, IDs: 3S84 and 1AV1). Secondary structure elements are highlighted (cyan, *α*-helix; purple, *β*-sheet; red, loop). **c** Correlations between docking scores (ipTM, interface ΔG) and structural features (peptide-region pLDDT, helix fraction of peptide or parent protein). The turquoise band shows the 95% confidence interval of the expected cumulative recovery of the top scoring candidates under random sampling. The blue line shows the observed cumulative recovery of the top scoring candidates. The top is defined as the peptides scoring in the last top quantile and arise almost exclusively from helical, high-confidence regions. **d** Randomly sampled encrypted peptides extracted from the same set of proteins identified in the wound fluid analysis were docked to CD14 or to PLY (PDB: 5AOD), and the calculated scores were correlated with structural features. **e** Randomly sampled encrypted peptides docked to CD14 (top row) show weaker correlations with helicity and pLDDT, without enrichment at the protein level. No correlations are detected for random peptides docked to PLY (bottom row).

To test whether this trend reflects a general property of AlphaFold, we performed control docking of randomly sampled encrypted peptides (**Fig. 5d**). When docked to CD14, these peptides showed a weaker but still detectable correlation between helicity, pLDDT and docking scores, whereas no enrichment was observed at the level of the parent protein. By contrast, peptides docked against the unrelated bacterial protein showed no such trends (**Fig. 5e**).

These comparisons suggest that CD14 can accommodate binding helical peptide segments, explaining the weak helicity trend in the CD14 random controls. However, the strong enrichment of helical, high-confidence binders and of helical source proteins was only observed when using peptides from endogenous wound fluid peptidome datasets. These findings indicate that the helix bias is not caused by prediction artifacts but is instead an intrinsic feature of the biological CD14-binding repertoire.

To improve our chances of finding binders to CD14 we filtered the results after docking the top 150 scoring binders with both AlphaFold2 and Boltz-2. In total, 12 peptides were identified to consistently dock with the same binding pose across models with low interface ΔG and high ipTM. Explicitly-solvated atomic-resolution molecular dynamics simulations over 200 ns showed that these candidates were stable over time and that binding free energies were negative for all 12 complexes, calculated with MMPBSA [40].

### Mining the complete human proteome for encrypted peptide binders

Mass spectrometry captures only a partial snapshot of the endogenous peptidome, whereas recent computational strategies have begun to interrogate complete proteomes for encrypted bioactive peptides [18,21,41]. Such approaches typically rely on exhaustive evaluation of all possible k-mers within the proteome or generating a predicted peptidome using heuristic rules such as proteasomal cleavage. While feasible for properties such as antimicrobial activity, exhaustive evaluation is impossible for structural binding assessment due to the computational load, rendering proteome-wide searches for peptide binders inaccessible to docking-based approaches.

We reasoned that the uncertainty-aware sampling strategy implemented in BoPep would enable exploration of complete proteomes by evaluating only a minute fraction of the available sequence space. To this end, we implemented an iterative sampling scheme in which peptides were drawn directly from the human proteome and evaluated iteratively using the BoPep framework. The search was initialized by docking an initial batch of randomly sampled peptides, which served to train the surrogate model. In subsequent iterations, large pools of candidate peptides were proposed to the surrogate model, ranked according to the acquisition function, and only the highest-ranked candidates were selected for docking and scoring. This procedure was repeated for a fixed number of iterations, allowing the search to progressively focus on informative regions of proteome-derived sequence space (**Fig. 7a**).

We applied this strategy to identify other candidate immunomodulatory peptides binding to CD14 using complete human proteome, excluding immunoglobulins, comprised of 20,357 proteins (UniProt download 13th of December 2025). In each iteration, 50,000 peptide candidates were sampled according to a uniform length distribution between 8-35, of which 10 were selected for docking. The search was seeded with 500 docked peptides, followed by 25 iterations prioritizing exploration through maximization of predictive uncertainty and 250 iterations prioritizing exploitation via expected improvement. In total, 13,750,000 (non-unique) peptides were evaluated by the model and 3,250 were docked.

The optimization dynamics differed from those observed in the peptidome-based search. Most notably, the surrogate model exhibited an immediate increase in predictive performance, with validation *R*^2^ values exceeding 0.7 (**Fig. 7b**). This phase coincided with the selection of peptides with consistently high objective values, suggesting exploitation of a local optimum within the proteome-derived search space (**Fig. 7c**). As the search progressed, model performance gradually decreased and converged to levels comparable to those observed in the peptidome search, consistent with exploration of more heterogeneous regions of sequence space.

Despite these differences, the overall behavior of the proteome search mirrored key features of the peptidome search. High-scoring peptides were progressively enriched across iterations (**Fig. 7c**), accompanied by systematic improvements in individual score components such as ipTM, peptide pAE, interface packing and interface ΔG (**Fig. 7d**). The highest-scoring peptide identified across both searches originated from glypican-1 and adopted a helical binding mode comparable to that observed for top-ranked peptides derived from the wound fluid peptidome (**Fig. 7c**).

To assess whether the proteome search accessed similar regions of sequence space as the peptidome search, the top-ranked proteome-derived peptides were projected onto the peptidome embedding space using UMAP. This analysis revealed an overlap between the regions occupied by high-scoring peptides from both searches (**Fig. 7e**), indicating that BoPep converges on shared structural and sequence features irrespective of whether candidates originate from experimentally observed peptidomes or from the complete proteome.

Together, these results demonstrate that BoPep enables tractable, structure-based exploration of entire proteomes for encrypted peptide binders. This expands the notion of proteome-wide searches beyond metrics like antimicrobial activity to metrics that are expensive to compute like peptide-protein binding, paving the way for future efforts in mining for encrypted peptide.

### *De novo* design of an antivirulence peptide capable of neutralizing pneumolysin

To further demonstrate the utility of BoPep, we evaluated the performance in combination with a generative design workflow, focusing on pneumolysin (PLY), a pore-forming toxin and a major virulence factor secreted by *Streptococcus pneumoniae* [31]. PLY exerts its effect by binding to cholesterol-rich host cell membranes followed by oligomerizing into large *β*-barrel pores, leading to cytolysis and the release of danger-associated molecular patterns. Neutralizing the effects of PLY is therefore advantageous, as reduced PLY activity will mitigate both bacterial virulence and host-driven immunopathology [42].

To generate inhibitors against PLY, we attached BoPep as a final filtering step after *de novo* peptide design using RFdiffusion and ProteinMPNN (**Fig. 8a**). The target region within PLY was a previously identified epitope that confers complete neutralization when targeted by a neutralizing monoclonal antibody (mAb). Binding site residues were defined by hotspot mapping of the neutralizing epitope identified through integrated structural modelling. Peptide backbone structures were generated using RFdiffusion and sequences were sampled using ProteinMPNN, followed by iterative cycles of Rosetta FastRelax and re-seeding, increasing sequence diversity. Each cycle generated new peptide candidates.

To establish how many designs to use as a starting point and how to determine the number of rounds of relaxation, a batch of 500 designs were relaxed for 20 iterations. Interface ΔG values improved only modestly after the first relaxation step, with subsequent refinements yielding diminishing returns (**Fig. 8b**,**c**). By the fourth round of MPNN and subsequent relaxation, structural diversity plateaued, indicating that four rounds were sufficient to capture most of the sequence-structure variability (**Fig. 8d**,**e**).

In the next step, BoPep was applied to search the generated sequence space, comprising 20,000 sequences across 230 iterations (5000 designs each undergoing four relaxation and MPNN rounds). In total 11.5% of search space was sampled to identify the top binder candidates (**Fig. 8f-h**). During the exploration phase, BoPep sampled broadly across the sequence space, before subsequently concentrating predictions around the two dominant structural families during exploitation (**Fig. 8i**). Although thousands of candidates were evaluated, the top designs converged into two dominant binding modes, a *β*-sheet-based pose and an *α*-helical pose, targeting different regions of the epitope (**Fig. 8j**,**k**).

To determine if these modes can bind and neutralize the cytolytic activity of PLY, we synthesized the 26 highest-scoring designs and tested if the synthesized peptides could compete out the high-affinity neutralizing mAb (Kd: 1.213 nM) and to inhibit PLY-mediated cytolysis (**Fig. 8l**). Using a competitive ELISA, we found that five peptides displaced the mAb out of which four formed *β*-sheets (**Fig. 8m**). Among these, two neutralized the cytolytic activity of PLY using a hemolysis inhibition assay. Both of the neutralizing peptides formed *β*-sheets when predicted to be interfacing with PLY (**Fig. 8n**).

Together, these findings show that BoPep is agnostic to the origin of the sequence space and can be readily integrated into binder discovery pipelines to identify bioactive protein-binding peptides while reducing computational screening effort. Using this workflow, we identified short peptides capable of displacing a high-affinity mAb and attenuating PLY-mediated hemolysis, thereby establishing a promising entry point for *de novo* design of antivirulence inhibitors targeting pneumolysin and potentially other bacterial virulence factors.

## Discussion

Peptides play a central role in biological research because they act as key signaling molecules and provide modular systems for probing protein activity. In addition, peptides represents a promising class of therapeutics, capable of selective protein binding while avoiding many of the challenges associated with large biologics. These properties make them comparatively easy to model and generate, while still providing the diversity needed to modulate specific protein–protein interactions. However, identifying functional binders within the vast peptide sequence space remains difficult, as exhaustive structural modelling or experimental screening represents a major bottleneck for both naturally occurring and *de novo* generated peptides.

In this study, we framed the task of identifying protein-binding peptides as an optimization problem in sequence space with an expensive black-box function, making Bayesian optimization (BO) a suitable strategy for efficient exploration. We approached this problem by developing a state-of-the-art BO pipeline that integrates recent developments in structural biology and deep learning. The pipeline consists of four main modules, including sequence embedding using protein language models, structural evaluation using structure prediction models, surrogate modelling using probabilistic recurrent deep learning architectures, and scoring using machine-learning informed metrics. Together, these components provide an adaptive solution for efficiently navigate the sequence space. This strategy is advantageous, as long as structural evaluations remain more expensive than surrogate updates, as is the case with docking or structural modelling.

The applications to CD14 and PLY demonstrate the utility of BoPep. In both cases, only a small fraction of the available sequences required evaluation before converging on peptides with favourable predicted interfaces, substantially reducing computational costs. These examples also demonstrate that BoPep can accommodate peptides derived from both *de novo* backbone generation, peptidomics data sets generated from endogenous proteolysis, and theoretical peptides encrypted in the human proteome. This flexibility enables exploration of both the encrypted peptide hypothesis by screening for bioactive peptides in biological and clinical samples, or in the proteome, and to be attached as a final step in generative pipelines to screen candidate binders. More broadly, the ability to effectively navigate large sequence landscapes positions BoPep as a tool for accelerating peptide discovery and for expanding the range of targetable protein–protein interaction.

The main limitations of BoPep stem for broader challenges in structural biology, most prominently related to inaccuracies in predicting binding poses and affinities, which directly affect the objective function BoPep optimizes for. Improvements in structural modelling and scoring, informed by larger experimental datasets, will likely enhance the objective, resulting in more efficient evaluations and higher success rates. Furthermore, advances in embeddings and surrogate architectures may offer similar gains, as BoPep is structured around a modular and problem-agnostic design so that new methods can be incorporated readily. This architecture also enables the use beyond peptide binder discovery, positioning it as a general tool for navigating large biological sequence spaces.

In summary, BoPep provides a principled and efficient approach for exploring peptide sequence spaces that would otherwise be intractable to explore exhaustively. By integrating structural scoring, uncertainty-aware modelling and Bayesian optimization, BoPep concentrates computational effort on the most informative regions of sequence space. This enables a practical and scalable strategy for identifying high-confidence peptide binders from both natural and synthetic peptide pools, including endogenous proteolytic fragments, full proteomes, and *de novo* design landscapes. As computational structure prediction, molecular docking, and protein–peptide interaction models continue to improve, optimization frameworks such as BoPep are likely to become increasingly important. Together, these features positions BoPep as a generalizable approach with relevance for the discovery and development of next-generation peptide-based therapeutics.

## Methods

### Overview of the BoPep workflow

BoPep is implemented in Python and requires inputs consisting of a target structure provided in PDB or CIF format, an optional binding site specification, and hyperparameters such as the surrogate model architecture, optimization schedule and dataset. The nature of the dataset depends on the search specification, as further described below. BoPep is modular at its core, with modules for embedding, co-folding, scoring, surrogate modelling, and main driver scripts for performing BO on a fixed peptidome, an entire proteome, and for designing peptides. When applying BoPep to search for peptide binders in a predefined search space, such as a peptidome, the input is a list of peptides. When applying the search to an entire proteome, the input is a dictionary of protein identifiers and sequences. In the design mode, the number of requested designs and the number of relaxation cycles must also be specified. Below we describe the individual components of BoPep, and lastly describe how these are strung together in search pipelines.

#### Initialization

Before the search begins, peptides are represented using either physicochemical characteristics from the AAindex database or embeddings from the protein language model ESM2 (650M). These feature spaces are high-dimensional and to improve learning over low-N datasets, dimensionality reduction is performed using either PCA or a VAE.

The search is initialized by co-folding an initial batch of peptide-target complexes using either the ColabFold implementation of AlphaFold2-multimer or Boltz-2. A template is provided for the target, whereas the peptide is predicted in single-sequence mode, removing the need for multiple sequence alignments. Initial peptide candidates are drawn either randomly or from the centroids of clusters obtained using k-nearest neighbors. The clustering strategy reduces redundancy by down-weighting dense regions in sequence space, thereby facilitating broader exploration.

#### Iterative search

At each iteration, protein-peptide complexes are scored using a combination of physicochemical features and structure-based confidence metrics. To allow optimization, these heterogeneous measures are combined into a single scalar objective, although BoPep also does support multiobjective optimization. The scalarization function can be provided by the user, but BoPep hosts a set of pre-defined functions we’ve developed. The search process aims to maximize or minimize this objective, which in the case for binder design ideally serves as a proxy for docking probability and pKd.

The surrogate model learns a mapping from peptide embeddings to the scalar objective. Three architectures are implemented: BiGRU, BiLSTM and MLP. In the MLP implementation we use stacked linear layers with ReLU activations. The recurrent network processes sequence embeddings with positional encodings that inject order information before a bidirectional GRU or LSTM. The recurrent hidden states are aggregated by a self-attention mechanism, complemented by masked max and mean pooling to handle variable sequence lengths, and passed to an MLP head. To capture predictive uncertainty, BoPep supports four probabilistic formulations which all output both a mean and a variance for each prediction.

For network ensembles, several networks are trained independently. Each network produces a prediction, and the mean and variance of these predictions are outputted. For MC dropout, a single network with dropout active at inference is run multiple times, producing stochastic predictions. The predictive mean and variance are computed in the same way as for the network ensemble.

For mean-variance estimation (MVE), the network directly outputs a mean and the log of the variance. Training is performed by minimizing the Gaussian negative log-likelihood which penalizes both large residual errors and unjustified variance inflation. This allows the network to model aleatoric uncertainty arising from noise in the training data.

For deep evidential regression (DER), the network outputs four parameters (*μ, v, α, β*) of a Normal-Inverse-Gamma prior over the predictive mean and variance. These are then used to compute a mean and variance. This parametrization allows us to split the variance into aleatoric and epistemic uncertainty. The training loss combines the negative log-likelihood of the distribution with a regularizer that penalizes evidence parameters when residuals are large [35]. This objective encourages the network to express uncertainty faithfully when data support is weak, while concentrating the predictive distribution when more evidence is available.

When using recurrent architectures, embeddings are represented as sequences of length *L* with reduced dimension *D*_reduced_, meaning that the embeddings are in domain 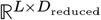. In the MLP case, embeddings are averaged to the domain 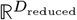.

The outputted mean and variances are used in the BO loop to compute acquisition values. BoPep supports different acquisition functions, that are used to rank candidates for selection.

Since the evaluated dataset grows dynamically during the search, surrogate hyperparameters are updated adaptively. In provided intervals, Bayesian hyperparameter optimization as implemented in Optuna is performed with the objective of minimizing the mean negative log-likelihood across crossvalidation folds.

#### Searching a pre-defined peptidome

When applying the search to a pre-defined peptidome, the user provides a list of peptides that constitute the search space, alongside a phase schedule which outlines the choice of acquisition function and number of iterations, as well as various search parameters. The search then follows the initialization and iterative search outlined above.

#### Searching a proteome

Applying the serach to a proteome requires a principled way of extracting peptides from the proteome. Here we apply a simple strategy in which peptides are randomly sampled from proteins according to a length distribution, by drawing a protein and position uniformly. The number of samples are specified by the user.

#### Designing peptides and searching among the generated peptides

For *de novo* design, the first step is to construct a large and diverse peptide-protein dataset to provide a search landscape for optimization. RFdiffusion is used to generate novel backbone structures by iterative denoising of Gaussian-perturbed backbone coordinates, followed by sequence recovery through cycles of ProteinMPNN and FastRelax. The generated sequence space is thereafter treated as a peptidome, and inputted into the pre-defined peptidome-search outlined above.

#### Modularity

The inherently modular build of BoPep allows it to be used for a variety of sequence and/or structure optimization problems. For example, instead of co-folding, one can fold without a target structure, and thereby optimizing for features of a single chain instead of a complex. Additionally, we have used its modularity to implement a generative model based on genetic algorithms and Bayesian methods not outlined here.

### Computational efficiency

The motivation behind BoPep lies in the computational cost of evaluating binders. This step is dominated by co-folding and scoring, and may also include alternative docking regimes or even molecular dynamics simulations. These evaluations require substantial wall time, even on costly GPUs, and any reduction in the number of docked candidates provides immediate computational benefit. In our applications to pre-defined peptidomes, BoPep identifies high-scoring peptides while evaluating roughly ten percent of all candidates. The additional computations introduced by the framework consist of embedding, surrogate model training and occasional hyperparameter optimization, whereas inference, score aggregation and acquisition computations contribute negligibly or are also present in standard workflows.

When using a pre-defined search space, the gain can be quantified by contrasting the exhaustive baseline, which requires *N t*_dock_ for *N* candidates and a per-docking cost *t*_dock_, with the BoPep runtime. Embedding imposes a fixed cost *T*_embed_, incurred once for the entire dataset. Surrogate training contributes a cost *T*_train_ at each iteration. For the small neural networks used here, a single training step typically requires about one minute on a modern GPU when optimizing over a batch of ten newly docked peptides. After *K* iterations, the total overhead becomes

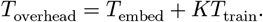

The full runtime is therefore

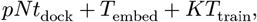

and BoPep is advantageous whenever

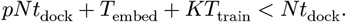

This condition is equivalent to

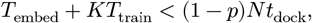

which shows that the method remains efficient provided that embedding and surrogate updates require less time than the block of avoided dockings.

Taking *t*_dock_ = 5 minutes means that in the CD14 mining run, we avoided (1−0.095)×35, 243 ≈ 31,900 dockings, corresponding to about 2,700 hours of single-machine wall time. Embedding the full dataset required less than one hour, and each of the *K* = 275 surrogate model updates added about 1-5 minutes. The total overhead therefore remained well below tens of hours, importantly far below the threshold set by the avoided dockings.

Finally, when searching across the complete proteome, the size of the candidate space becomes prohibitive. For a set of peptide lengths (e.g. *k* = 8–35), the total number of k-mers is

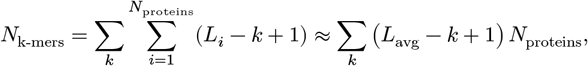

which evaluates to approximately 2.1 × 10^8^ candidates for *L*_avg_ = 400 and 20,000 proteins. Even with relatively inexpensive evaluation functions, such a search space is effectively intractable. In this regime, BoPep acts as an optimizer over a predefined yet near-infinite combinatorial space, enabling targeted exploration that would otherwise be computationally infeasible.

### Creating a predictive equation for ranking binders

To construct an objective function that can act as a ranking proxy for binding probability and affinity, we curated a set of linear peptide-protein complexes with associated dissociation constants from PDBbind [38] (downloaded April 3rd 2025). Structures were filtered to retain entries comprising a single receptor chain and a single peptide chain composed of canonical amino acids, without additional small-molecule ligands. Proteins were stripped of peptide coordinates and used as targets for docking, in which each peptide sequence was re-docked and scored within BoPep. Complexes that did not recover the native pocket were excluded. The resulting set consisted of predicted complexes that aligned to the original pose and for which experimental affinity could be expressed as pKd.

To quantify specificity, we constructed matched decoy cohorts by two procedures. First, each native peptide was residue-shuffled to preserve composition while disrupting sequence order. Second, independent random peptides were sampled according to the naturally occurring amino acid distribution. All decoys were co-folded against their corresponding targets and scored using the same BoPep pipeline. Real and decoy data were partitioned into training, validation and test splits.

From each predicted complex we computed scores using the BoPep scoring framework. These features formed the input to two supervised tasks. The first was a binary classifier trained to distinguish native complexes from decoys. The second was a regressor trained on native complexes to predict pKd. We fitted baseline models including logistic or linear regression, random forests and support vector machines.

In parallel, we performed symbolic regression to derive low-complexity, closed-form expressions for both tasks using PySR [37]. The search proceeded by evolutionary operations over an operator set comprising addition, subtraction, multiplication, division and affine transformations. The search optimized for L2-loss and model complexity. For the classifier, the symbolic score was converted to a class label by applying a threshold of 0.5. For the regressor, expressions were evaluated by the correlation to experimental pKd.

The classification term was defined as

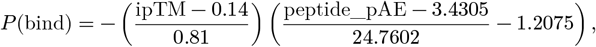

and the regression term as:

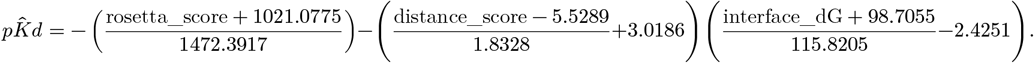

The final BoPep objective was then obtained by multiplying 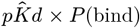. Peptides that are not predicted to dock in the binding site, defined by proximity with a list of residues, are penalized, and their objective value is set to −1. This construction retains the discriminative power of the classifier while scaling predictions by a quantitative affinity trend in the binding site. After selection, the composite expression was locked and evaluated once on the held-out test set.

### Benchmarking embeddings, surrogate architectures and uncertainty estimation

To evaluate the impact of different embedding strategies, surrogate architectures, and uncertainty estimation modalities on optimization performance, we implemented a controlled benchmarking protocol. For all benchmarks, we used data from the endogenous wound fluid peptidome dataset (see *Mining encrypted immunomodulatory binders to CD14* for details).

To investigate the embedding methods, peptides were embedded either using amino acid physicochemical indices from the AAindex database or with the ESM2-650M protein language model. Dimensionality reduction was applied using PCA or VAEs. We assessed local consistency and global organization using UMAP projections.

To asess the performance during BO, 500 random peptides were selected from the dataset. These served as a fixed search landscape on which BO was repeatedly applied under varying configurations, with maximizing ipTM as the reward objective. Each surrogate model modality and architecture and hyperparameter was tuned separately for learning rate, hidden size, and uncertainty parameters (number of ensembles, dropout ratio, and tuning parameters for MVE and DER) using cross-validation. All benchmarks were run in triplicate. For each configuration, BO was executed for 40 iterations with expected improvement acquisition. The search was initialized with 50 complexes. A test set of 50 complexes was used to evaluate surrogate model performance. The batch size was set to 10.

### Mining the peptidome for encrypted immunomodulatory binders to CD14

A mass-spectrometry derived peptidomic dataset was curated by integrating data from clinical sources. These include the wound fluid peptidomes from: non-healing leg ulcers (N=18, four patients, PXD: PXD048892), infected wounds (N=3, datamatrices available in [14]), and acute surgical drainage (N=5, datamatrices available in [14]).

The structure of CD14 used here is based on the crystal structure deposited to the PDB (PDB: 4GLP). Similarly to in Petruk et al. [11], 5 residues were added to the N-terminal (residues 20-25) alongside a disulfide bridge (C25-C36) since these residues are proximate to the binding site and may affect predictions.

A filtering step was performed to remove peptides with repetitive sequences or other characteristics that were unlikely to result in strong binding interactions and that introduced artificially high variations in the data, potentially misguiding the optimization process. Peptides with overly repetitive sequences, such as those consisting of a single amino acid repeated multiple times were excluded from further consideration. The final dataset consisted of 35,243 peptide sequences. The target residues were set as 22, 23, 24, 42, 43, 44, 45, 46, 47, 48, 49, 50, 51, 52, 53, 69, 70, 71, 72, 73, 74, 75, 76, 77, 81, 82, 83, 84, 85, 86, 87, 88, 89, 90, 104, 105, 106, 107, 108, 109, 110. The batch size was set to 10. We found that 8 contacts (defined as residues within <5 Å) reliably selected for peptides in the desired pocket. The validation split was set to 0.2. Hyperparameter optimization was performed every 25 iterations with 20 trials and 3 splits in the cross-evaluation.

BoPep was initialized with 500 dockings, and used to search the peptidome. The acquisition function in the first 25 iterations aimed to select peptides with the highest variance. This was followed by 250 iterations of expected improvement acquisition. For the final 10 iterations, peptides with the highest predicted mean scores were selected. This resulted in a total of 3350 dockings, including the seeding 500 initial ones. The number of AlphaFold models was set to 5 and the number of recyles to 10, with a early stop tolerance of 0.1 Å.

### Analyzing helical propensity among CD14 binders

We examined whether BoPep preferentially selects peptides from high-confident *α*-helical regions. For each peptide evaluated in the CD14 search, helix fraction was estimated from sequence composition and from DSSP assignments of AlphaFold monomer models. These were downloaded using the Alphafold server API. Per-residue pLDDT was extracted and averaged across the peptide span and across the parent protein. These features were correlated with docking scores (ipTM, interface ΔG, composite objective).

To test for preferential selection, peptides were ranked by each structural feature and the cumulative number of top-scoring peptides was computed along the ranked list. This empirical enrichment was compared to a null expectation assuming that high-scoring peptides are randomly distributed with respect to the feature The null distribution of cumulative counts was modeled using a hypergeometric distribution, reflecting sampling without replacement from a finite peptide pool, and used to derive confidence bands for the random expectation.

A subset of 200 complexes was also docked with Boltz-2 to confirm that similar associations were obtained. To test for model-intrinsic bias, we generated control sets of 300 random encrypted peptides by sampling spans of 8-30 residues uniformly from AlphaFold-predicted proteins. These peptides were docked either to CD14 or to PLY. For each peptide, helicity and pLDDT were computed as above and compared with docking metrics.

### Molecular dynamics simulations and binding free energy computations

To mitigate the impact of biases and achieve higher confidence results the peptides were filtered to those with consistent high quality predictions from both AF2 and Boltz-2. This resulted in 12 peptides (**Table 1**).

**Table 1.**
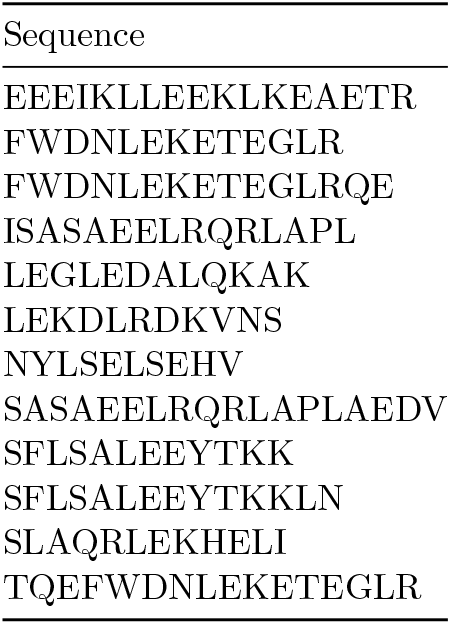
Top scoring peptides docked against CD14.

These were subjected to atomic resolution molecular dynamics (MD) simulations. To reduce computational resources required to run these simulations, we truncated residues 131-335 in CD14 and restrained the C-terminus as extensive microsecond timescale simulations with the binder sHVF18 showed similar peptide binding profile and stability to the full-length protein simulations performed in our previous study (**Fig. 6**) [11].

**Fig. 6.**
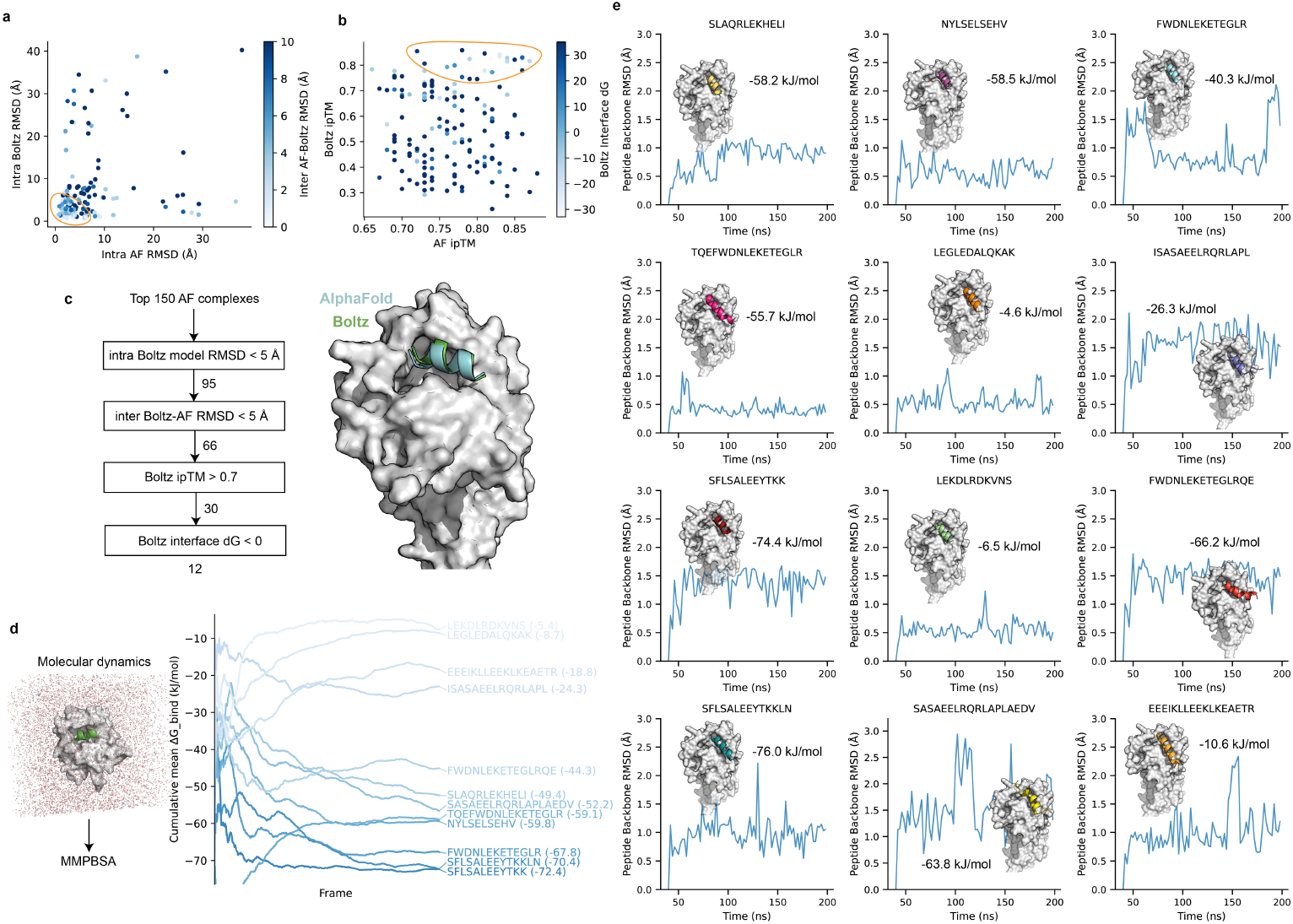
Molecular dynamics identify stable peptide-protein complexes. **a** Comparison of peptide backbone RMSDs between Boltz-derived models and AF-derived models of the top 150 scoring peptide candidates identified in the search. The color scale indicates the model peptide backbone root mean squared error (RMSD, Å) between both AF and Boltz. **b** Relationship between AF ipTM and Boltz ipTM, colored by Boltz interface binding free energy (ΔG). **c** Schematic illustration of the filtering workflow for selecting high-confidence peptide-protein complexes. From the top 150 AF models, sequential filtered by intra-model agreement, inter-method agreement, ipTM, and interface ΔG resulting in 12 top-ranking candidates. Representative structure shows alignment between the best AF (grey) and Boltz (green) models for a representative peptide. **d** The 12 candidates were subject to 200 ns explicitly-solvated atomic-resolution molecular dynamics simulation, and binding energies were estimated with MMPBSA. The plot shows the mean cumulative ΔG across the frames. The annotated energies are the final estimates. **e** Peptide RMSD of the molecular dynamics trajectories for the 12 peptides, annotated with the binding free energy and a snapshot of the trajectory.

**Fig. 7.**
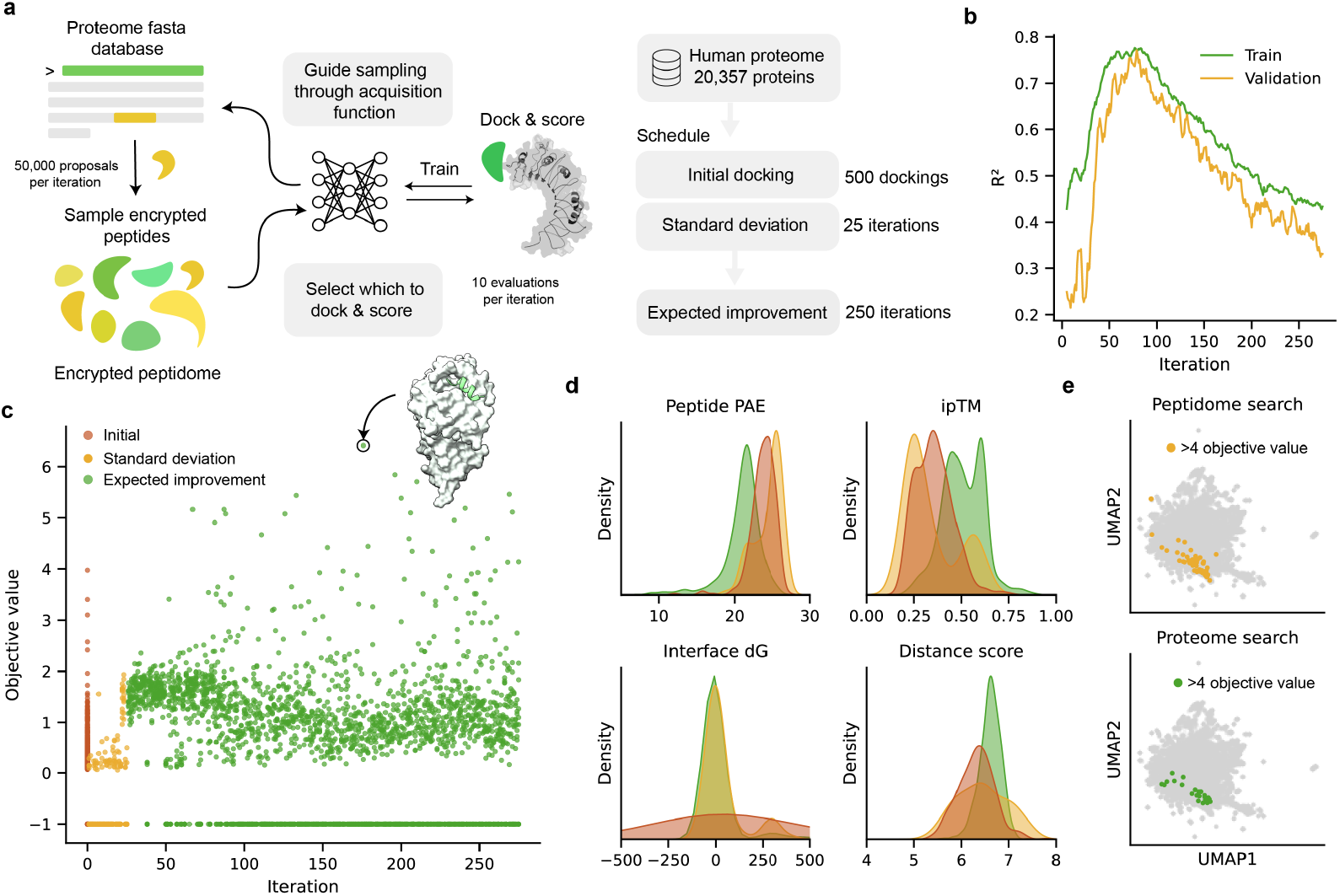
BoPep enables proteome-wide discovery of encrypted peptide binders. **a** Schematic of the iterative BoPep workflow applied to the complete human proteome. Peptide candidates are sampled directly from a database of 20,357 human proteins. In each iteration, large candidate pools of 50,000 peptides were evaluated by a surrogate model, ranked by an acquisition function, and a small subset of 10 peptides was selected for docking and scoring against the target protein. Docking results are fed back to retrain the model, enabling uncertainty-aware exploration and exploitation of proteome-derived sequence space. **b** Predictive performance of the surrogate model during the proteome search, shown as training and validation *R*^2^ over iterations. **c** Objective values of docked peptides across iterations, colored by acquisition strategy. The inset shows the predicted complex of the top scoring peptide, KLVSEAKAQLRDV derived from glypican-1.**d** Distributions of individual score components for docked peptides, including peptide pAE, ipTM, interface ΔG (ΔG), and distance score. **e** UMAP projection of the wound fluid peptidome and the evaluated peptide candidates from the proteome search. The peptides with objective values >4 identified during the peptidome and proteome search are highlighted in the top and bottom plot respectively.

**Fig. 7.**
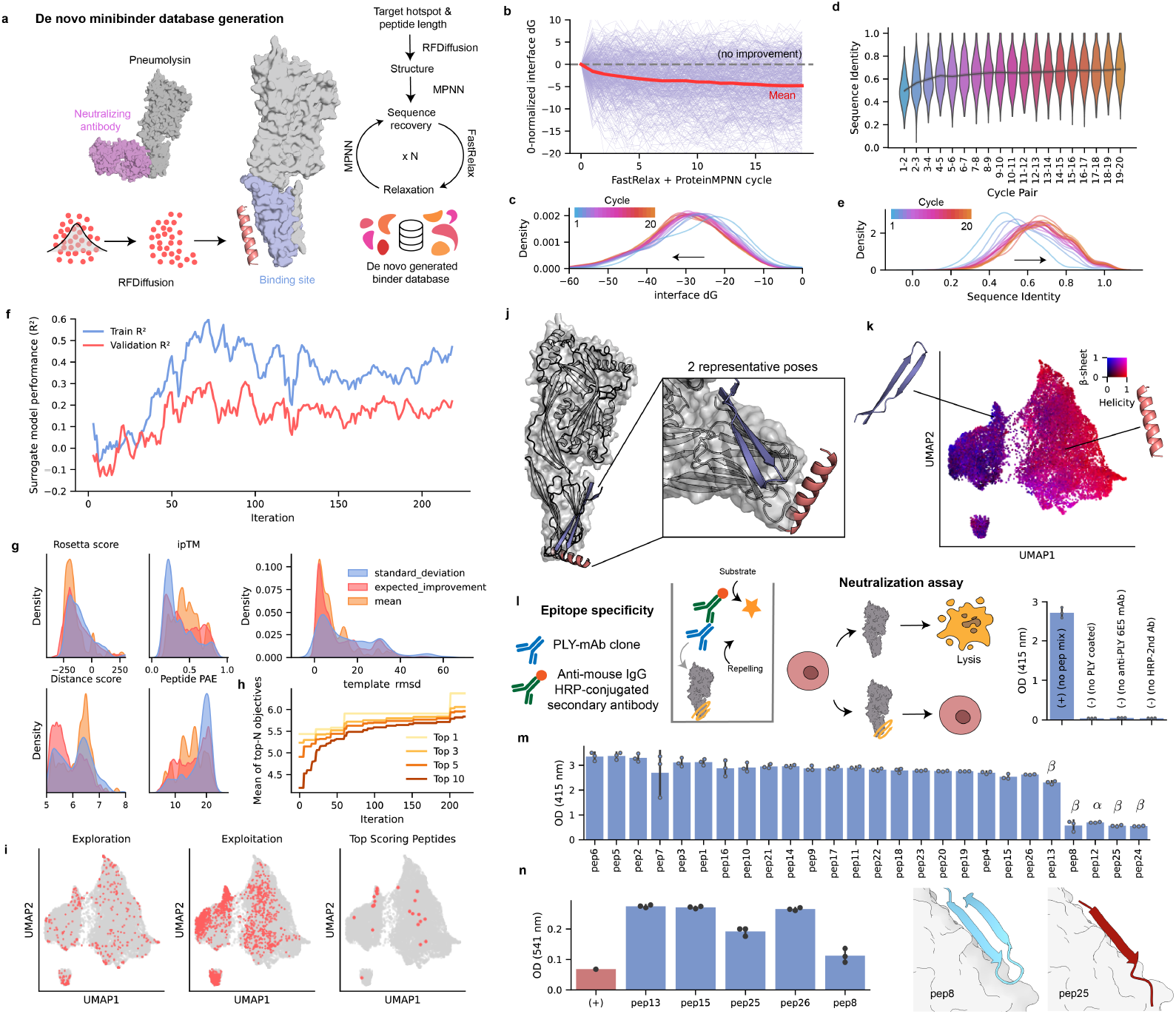
BoPep enables effective screening of *de novo* binder design against PLY. **a** Schematic overview of workflow to generate peptide binders to PLY. Binding site residues were defined using information from a previously identified epitope targeted by a neutralizing antibody. RFdiffusion was used to design peptide backbones which were then subjected to sequence recovery with protein MPNN, iterative relaxation, and evaluation, producing a peptide database consisting of 20,000 sequences. **b** Lineage plot of the interface energy (ΔG) after iterative FastRelax + MPNN cycles. **c** Kernel density estimate plots of interface ΔG across cycles. **d**,**e** Pairwise sequence diversity across design cycles, shown as violin plots (**d**) and pairwise sequence identity distributions (**e**). **f** Line plot showing the performance of the surrogate model over optimization iterations for training and validation sets. **g** Distribution of Rosetta interface scores, ipTM scores, distance errors, peptide pAE values, and template RMSD during optimization, colored by acquisition function (mean, standard deviation, expected improvement). **h** The mean objective value of the top-ranked peptides across iterations. **i** UMAP projections of the embeddings of the designed peptides, highlighting exploration (left), exploitation (middle), and clustering of top-scoring peptides (right). **j** Structural models of two representative peptides bound to PLY. **k** UMAP projection of the embeddings of the designed peptides, colored by predicted helicity/beta-sheet propensity. **l** Schematic of the experimental validation workflow. Designed peptides were evaluated using a competitive ELISA in which peptides were incubated with PLY in the presence of the neutralizing monoclonal antibody 6E5 and a secondary antibody conjugated with HRP. Absence of any peptide serves as the positive control; while no PLY, no antibody 6E5 or no secondary antibody served as negative controls. The selected peptides were subjected to PLY-neutralization assay to quantify the degree of inhibition. The right panel shows the OD_415_ values for the competitive ELISA assay controls. **m** Competitive ELISA of the 26 synthesized peptides. Bars show OD_415_ measurements from peptide–PLY–6E5 incubations, benchmarked against the baseline and negative controls shown in **l**. Lower OD reflects reduced binding of 6E5 in the presence of a peptide. Peptides annotated *α* or *β* indicate the predicted structural class of their dominant binding pose based on the docking models in **j**,**k. n** PLY-haemolysis inhibition assay used to assess the ability of designed peptides to inhibit PLY-mediated haemolysis. OD_541_ values represent the amount of released haemoglobin after exposing red blood cells to PLY with or without peptides, referenced to the assay controls (no-PLY as positive control). Representative structural models of pep8 and pep25 at their predicted PLY interfaces are shown. Error bars represent mean ± SD from three independent experiments.

The peptide-truncated CD14 complexes were parametrized using the CHARMM36m forcefield [43]. The complexes were solvated with TIP3P water molecules and 0.15 M NaCl salt was subsequently added to neutralize the system. Steepest descent energy minimization was then performed to remove any atomic clashes. A short 125 ps equilibration was performed with positional restraints (force constant, Fc = 400 kJ mol^−1^nm^−2^) applied to all backbone atoms of the protein and peptide. Afterwards a 200 ns production MD simulations was performed for each complex using GROMACS 2024 [44]. To maintain the stability of the truncated CD14 during the production runs, we applied positional restraints (Fc = 500 kJ mol^−1^nm^−2^) to the backbone atoms of residues 120-130. The temperature of the system was maintained at 310 K using the Nosé-Hoover thermostat [45,46], while the pressure was maintained at 1 atm by an isotropic coupling to a Parrinello-Rahman barostat [47]. Electrostatic interactions were computed using the particle mesh Ewald (PME) method [48], while the van der Waals interactions were truncated at 1.2 nm with a force switch function applied between 1.0-1.2 nm. A 2-fs integration timestep was used whereby all covalent bonds involving hydrogen atoms were constrained using the LINCS algorithm [49].

The binding free energy of the complexes was estimated using the molecular mechanics Poisson-Boltzmann surface area (MMPBSA) approach [40] as implemented in gmx_MMPBSA [50]. The binding free energy is computed as

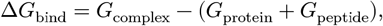

by `gmx_MMPBSA`, where each free energy term was decomposed into molecular mechanics (MM) energy in vacuum, polar solvation energy from the Poisson-Boltzmann (PB) model, and nonpolar solvation energy estimated from solvent-accessible surface area (SASA).

PB calculations were performed with an internal dielectric constant of 4.0 and a solvent dielectric constant of 78.5 to mimic water at 310 K. The ionic strength was set to 0.15 M, consistent with the simulation conditions. The solvent probe radius was 1.4 Å, with grid spacing defined by a scale factor of 2.0 and a fill ratio of 4.0.

### Mining the proteome for encrypted immunomodulatory binders to CD14

The human proteome was downloaded from UniProt (on 13th of December 2025). Immunoglobulins were removed from the database by removing all proteins containing “immunoglobulin” in their description. The same CD14 model was used as when identifying binders in the wound fluid peptidome. The search was seeded by sampling 500 peptides from the peptidome. All sampling of peptides was conducted by first sampling a random protein (uniformly) and then sampling a midpoint in that protein (uniformly) whereby the peptide was extracted by taking the amino acid sequence around the midpoint of length *L* ∼ *U*(8, 35). Peptides were that didn’t pass the filter described above were discarded.

The search used the same docking, objective, and binding site parameters as in the wound fluid search. Each iteration, 50,000 peptides were sampled from the proteome. Sequences were embedded with ESM, reduced by PCA, and the search was guided by a BiGRU DER surrogate. Hyperparamter optimization was conducted every 50th iteration. The acquisition function in the first 25 iterations aimed to select peptides with the highest variance. This was followed by 250 iterations of expected improvement acquisition, with a batch size of 10, totaling 3250 dockings.

### *De novo* peptide design against pneumolysin

Peptide backbones were generated with RFdiffusion, using the beta-sheet checkpoint to promote structural diversity. Binding hotspots were sampled from residues 370-460. The number of hotspots were samples from 𝒩(4, 2) truncated at [1, 8]. Peptide lengths were drawn from a uniform distribution *U*(10, 30). Sequences were recovered with ProteinMPNN, followed by iterative cycles of Rosetta FastRelax and ProteinMPNN to broaden sequence diversity while relieving local strain.

A pilot set of 500 designs was subjected to 20 FastRelax-ProteinMPNN cycles to determine an efficient schedule. Interface ΔG improved primarily after the first relaxation and plateaued by the fourth cycle, with a concurrent saturation of structural diversity. Guided by these observations, the full design campaign generated 5,000 RFdiffusion backbones and applied four relaxation-recovery cycles, yielding 20,000 unique sequences.

This database served as the search landscape for BoPep. Sequences were embedded with ESM, reduced by PCA, and the search was guided by a BiGRU DER surrogate. 100 peptides were used to seed the search. The acquisition schedule followed 30 iterations of standard deviation, 180 iterations of expected improvement, and 10 iterations maximizing the mean objective, resulting in 2300 dockings. The required number of contacts was set to 5. Batch size was set to 10. Hyperparameters were optimized every 50 iterations with 20 trials and 3 splits during cross-validation. The number of AlphaFold models was set to 5 and the number of recyles to 10, with a early stop tolerance of 0.1 Å. 26 top scoring peptides were selected for downstream analyses (**Table 2**).

**Table 2.**
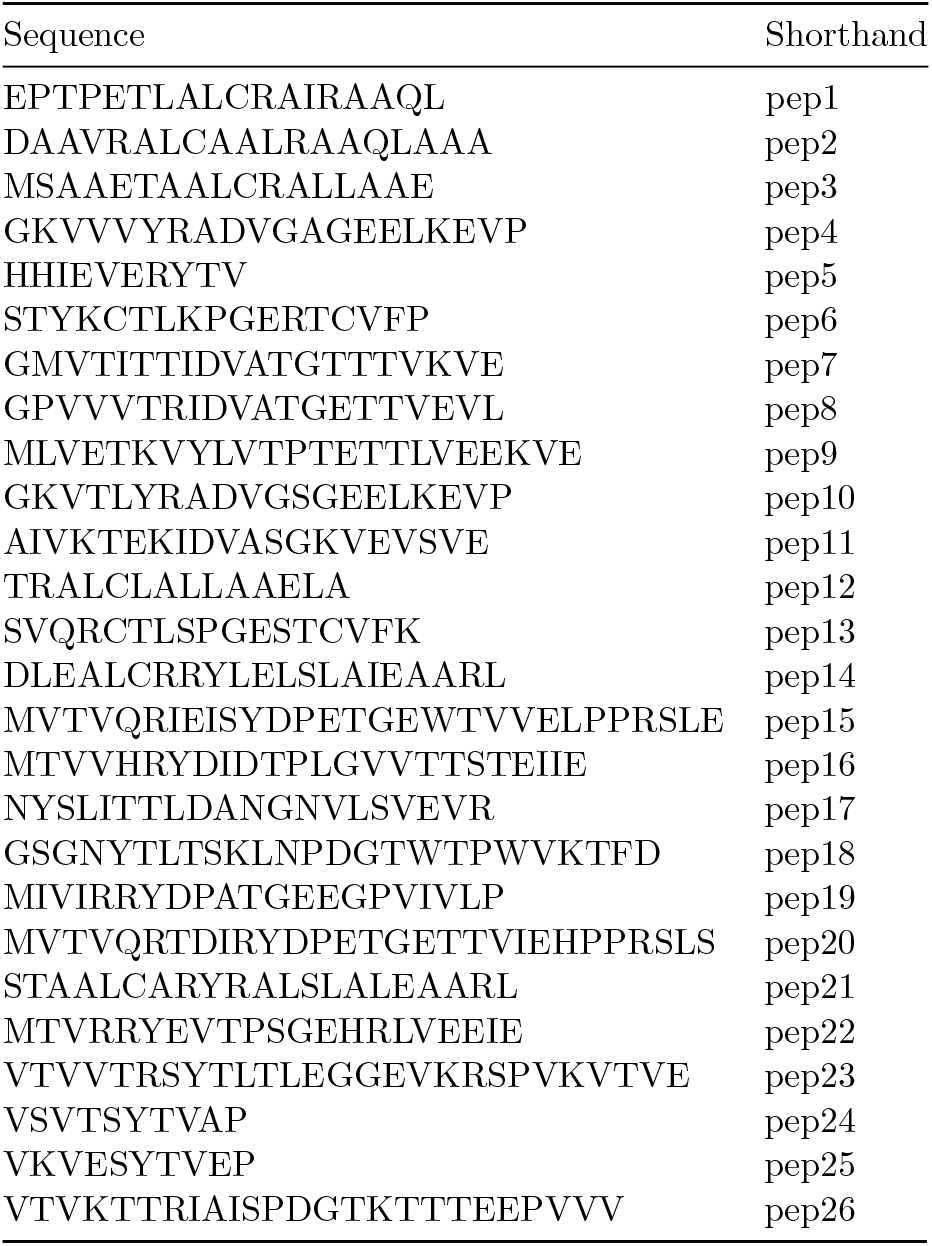
Top scoring designed sequences against PLY and that were synthesized and tested.

### Competitive ELISA and pneumolysin hemolysis inhibition assay

To evaluate peptide specific binding to the target PLY, competitive enzyme-linked immunosorbent assays (ELISAs) were performed. Equal amounts of purified PLY antigen were immobilized onto MaxiSorp 96-well plates (Thermo Fisher Scientific) and incubated overnight at 4 °C. Plates were washed three times with PBST (phosphate-buffered saline containing 0.05% Tween-20, Fisher Scientific) between all steps. Non-specific binding sites were blocked with freshly prepared 2% (w/v) bovine serum albumin (BSA, Sigma-Aldrich) for 1 h at room temperature (RT). Next, each designed peptide was added at 10 µg per well, and all peptide-PLY combinations were tested in triplicate. Following incubation for 90 min at RT with gentle agitation (300 rpm, ThermoMixer, Eppendorf), a predefined subnanomolar-affinity anti-PLY monoclonal antibody was introduced and incubated for an additional 35 min under the same conditions. Subsequently, plates were incubated sequentially with a horseradish peroxidase (HRP)-conjugated secondary antibody (1:3000 dilution) and a freshly prepared HRP substrate solution (Bio-Rad). After 5 min of color development, reactions were stopped by the addition of 100 mM H2SO4. Absorbance was measured at 415 nm wavelength using a microplate reader (BMG LABTECH). Competitive inhibition (or repelling) of PLY-antibody binding by each peptide was determined as the change in absorbance value relative to control wells where no peptide was added.

The capacity of peptides, displaying various repelling effect, to inhibit PLY-induced hemolysis was assessed as previously described. Briefly, 40 ng of recombinant active PLY toxin was incubated with either a neutralizing monoclonal antibody (mAb) or peptides in a two-fold serial dilution, starting from 2 µg (antibody) or 4 µg (peptide) per well. The mixtures were incubated in round-bottom 96-well plates (Corning), at 37 °C for 30 min to allow peptide/mAb-PLY complex formation. Next, freshly washed (2x) and diluted (4% v/v) sheep erythrocyte suspensions (Fisher Scientific) were added to each condition and incubated for an additional 60 min at 37 °C. Following incubation, plates were centrifuged at 500x g for 8 min, and the supernatants were carefully transferred to flat-bottom 96well plates (Corning) for absorbance measurement at 541 nm wavelength. The extent of hemolysis inhibition was calculated as the reduction in OD in comparisons. The applied neutralizing anti-PLY mAb served as the positive control, whereas the peptide solvent served as the vehicle control. All peptide-PLY conditions were tested in triplicate setting.

## Code and data availability

BoPep is available at GitHub under an MIT license. Binding data is available at PDBbind (https://www.pdbbind-plus.org.cn/).

## Acknowledgements

We thank Jonas Wallin for advice on probabilistic neural networks and Bayesian optimization, and Lars Malmström for guidance on the usage of MMPBSA.

## References

1. Muttenthaler M, King GF, Adams DJ, Alewood PF. Trends in peptide drug discovery. Nature Reviews Drug Discovery [Internet]. 2021 Feb;20[4]:309–25. Available from: 10.1038/s41573-020-00135-8

2. Capecchi A, Reymond JL. Peptides in chemical space. Medicine in Drug Discovery [Internet]. 2021 Mar;9:100081. Available from: 10.1016/j.medidd.2021.100081

3. Hellinger R, Sigurdsson A, Wu W, Romanova EV, Li L, Sweedler JV, et al. Peptidomics. Nature Reviews Methods Primers [Internet]. 2023 Mar;3[1]. Available from: 10.1038/s43586-023-00205-2

4. Hartman E, Forsberg F, Kjellström S, Petrlova J, Luo C, Scott A, et al. Peptide clustering enhances large-scale analyses and reveals proteolytic signatures in mass spectrometry data. Nature Communications [Internet]. 2024 Aug;15[1]. Available from: 10.1038/s41467-024-51589-y

5. Secher A, Kelstrup CD, Conde-Frieboes KW, Pyke C, Raun K, Wulff BS, et al. Analytic framework for peptidomics applied to large-scale neuropeptide identification. Nature Communications [Internet]. 2016 May;7[1]. Available from: 10.1038/ncomms11436

6. Foreman RE, George AL, Reimann F, Gribble FM, Kay RG. Peptidomics: A review of clinical applications and methodologies. Journal of Proteome Research [Internet]. 2021 Jul;20[8]:3782– 97. Available from: 10.1021/acs.jproteome.1c00295

7. Zasloff M. Antimicrobial peptides of multicellular organisms. Nature [Internet]. 2002 Jan;415[6870]:389–95. Available from: 10.1038/415389a

8. Ganz T. Defensins: Antimicrobial peptides of innate immunity. Nature Reviews Immunology [Internet]. 2003 Sep;3[9]:710–20. Available from: 10.1038/nri1180

9. Mookherjee N, Hancock REW. Cationic host defence peptides: Innate immune regulatory peptides as a novel approach for treating infections. Cellular and Molecular Life Sciences [Internet]. 2007 Feb;64[7–8]:922–33. Available from: 10.1007/s00018-007-6475-6

10. Oliveira Júnior NG, Souza CM, Buccini DF, Cardoso MH, Franco OL. Antimicrobial peptides: Structure, functions and translational applications. Nature Reviews Microbiology [Internet]. 2025 Jul;23[11]:687–700. Available from: 10.1038/s41579-025-01200-y

11. Petruk G, Puthia M, Samsudin F, Petrlova J, Olm F, Mittendorfer M, et al. Targeting toll-like receptor-driven systemic inflammation by engineering an innate structural fold into drugs. Nature Communications [Internet]. 2023 Sep;14[1]. Available from: 10.1038/s41467-023-41702-y

12. Nordahl EA, Rydengård V, Nyberg P, Nitsche DP, Mörgelin M, Malmsten M, et al. Activation of the complement system generates antibacterial peptides. Proceedings of the National Academy of Sciences [Internet]. 2004 Nov;101[48]:16879–84. Available from: 10.1073/pnas.0406678101

13. Papareddy P, Rydengård V, Pasupuleti M, Walse B, Mörgelin M, Chalupka A, et al. Proteolysis of human thrombin generates novel host defense peptides. Isberg RR, editor. PLoS Pathogens [Internet]. 2010 Apr;6[4]:e1000857. Available from: 10.1371/journal.ppat.1000857

14. Hartman E, Wallblom K, Plas MJA van der, Petrlova J, Cai J, Saleh K, et al. Bioinformatic analysis of the wound peptidome reveals potential biomarkers and antimicrobial peptides. Frontiers in Immunology [Internet]. 2021 Feb;11. Available from: 10.3389/fimmu.2020.620707

15. Nordahl EA, Rydengård V, Mörgelin M, Schmidtchen A. Domain 5 of high molecular weight kininogen is antibacterial. Journal of Biological Chemistry [Internet]. 2005 Oct;280[41]:34832–9. Available from: 10.1074/jbc.M507249200

16. Petrlova J, Hansen FC, Plas MJA van der, Huber RG, Mörgelin M, Malmsten M, et al. Aggregation of thrombin-derived c-terminal fragments as a previously undisclosed host defense mechanism. Proceedings of the National Academy of Sciences [Internet]. 2017 May;114[21]. Available from: 10.1073/pnas.1619609114

17. Pasupuleti M, Schmidtchen A, Malmsten M. Antimicrobial peptides: Key components of the innate immune system. Critical Reviews in Biotechnology [Internet]. 2011 Nov;32[2]:143–71. Available from: 10.3109/07388551.2011.594423

18. Goldberg K, Lobov A, Antonello P, Shmueli MD, Yakir I, Weizman T, et al. Cell-autonomous innate immunity by proteasome-derived defence peptides. Nature [Internet]. 2025 Mar;639[8056]:1032–41. Available from: 10.1038/s41586-025-08615-w

19. Kasetty G, Papareddy P, Kalle M, Rydengård V, Walse B, Svensson B, et al. The c-terminal sequence of several human serine proteases encodes host defense functions. Journal of Innate Immunity [Internet]. 2011;3[5]:471–82. Available from: 10.1159/000327016

20. Kalle M, Papareddy P, Kasetty G, Tollefsen DM, Malmsten M, Mörgelin M, et al. Proteolytic activation transforms heparin cofactor II into a host defense molecule. The Journal of Immunology [Internet]. 2013 Jun;190[12]:6303–10. Available from: 10.4049/jimmunol.1203030

21. Torres MDT, Cesaro A, Fuente-Nunez C de la. Peptides from non-immune proteins target infections through antimicrobial and immunomodulatory properties. Trends in Biotechnology [Internet]. 2025 Jan;43[1]:184–205. Available from: 10.1016/j.tibtech.2024.09.008

22. Lin Z, Akin H, Rao R, Hie B, Zhu Z, Lu W, et al. Language models of protein sequences at the scale of evolution enable accurate structure prediction. bioRxiv. 2022;

23. Jumper J, Evans R, Pritzel A, Green T, Figurnov M, Ronneberger O, et al. Highly accurate protein structure prediction with AlphaFold. Nature [Internet]. 2021 Jul;596[7873]:583–9. Available from: 10.1038/s41586-021-03819-2

24. Pacesa M, Nickel L, Schellhaas C, Schmidt J, Pyatova E, Kissling L, et al. One-shot design of functional protein binders with BindCraft. Nature [Internet]. 2025 Aug;646[8084]:483–92. Available from: 10.1038/s41586-025-09429-6

25. Lin Z, Akin H, Rao R, Hie B, Zhu Z, Lu W, et al. Evolutionary-scale prediction of atomic-level protein structure with a language model. Science [Internet]. 2023 Mar;379[6637]:1123–30. Available from: 10.1126/science.ade2574

26. Passaro S, Corso G, Wohlwend J, Reveiz M, Thaler S, Somnath VR, et al. Boltz-2: Towards accurate and efficient binding affinity prediction. bioRxiv. 2025;

27. Watson JL, Juergens D, Bennett NR, Trippe BL, Yim J, Eisenach HE, et al. De novo design of protein structure and function with RFdiffusion. Nature [Internet]. 2023 Jul;620[7976]:1089– 100. Available from: 10.1038/s41586-023-06415-8

28. Stark H, Faltings F, Choi M, Xie Y, Hur E, O’Donnell T, et al. BoltzGen: Toward universal binder design. https://hannes-stark.com/assets/boltzgen.pdf; 2025.

29. Li Q, Vlachos EN, Bryant P. Design of linear and cyclic peptide binders of different lengths from protein sequence information. 2024 Jun; Available from: 10.1101/2024.06.20.599739

30. Rettie SA, Juergens D, Adebomi V, Bueso YF, Zhao Q, Leveille AN, et al. Accurate de novo design of high-affinity protein-binding macrocycles using deep learning. Nature Chemical Biology [Internet]. 2025 Jun; Available from: 10.1038/s41589-025-01929-w

31. Pereira JM, Xu S, Leong JM, Sousa S. The yin and yang of pneumolysin during pneumococcal infection. Frontiers in Immunology [Internet]. 2022 Apr;13. Available from: 10.3389/fimmu.2022.878244

32. Kawashima S, Ogata H, Kanehisa M. AAindex: Amino acid index database. Nucleic Acids Research [Internet]. 1999 Jan;27[1]:368–9. Available from: 10.1093/nar/27.1.368

33. Mirdita M, Schütze K, Moriwaki Y, Heo L, Ovchinnikov S, Steinegger M. ColabFold: Making protein folding accessible to all. Nature Methods [Internet]. 2022 May;19[6]:679–82. Available from: 10.1038/s41592-022-01488-1

34. Nix DA, Weigend AS. Estimating the mean and variance of the target probability distribution. Proceedings of 1994 IEEE International Conference on Neural Networks (ICNN’94) [Internet]. 1994;1:55–60 vol.1. Available from: https://api.semanticscholar.org/CorpusID:117583961

35. Amini A, Schwarting W, Soleimany A, Rus D. Deep evidential regression. 2019; Available from: https://arxiv.org/abs/1910.02600

36. Leman JK, Weitzner BD, Lewis SM, Adolf-Bryfogle J, Alam N, Alford RF, et al. Macromolecular modeling and design in rosetta: Recent methods and frameworks. Nature Methods [Internet]. 2020 Jun;17[7]:665–80. Available from: 10.1038/s41592-020-0848-2

37. Cranmer M. Interpretable machine learning for science with PySR and SymbolicRegression.jl [Internet]. arXiv; 2023. Available from: https://arxiv.org/abs/2305.01582

38. Wang R, Fang X, Lu Y, Yang CY, Wang S. The PDBbind database: methodologies and updates. Journal of Medicinal Chemistry [Internet]. 2005 May;48[12]:4111–9. Available from: 10.1021/jm048957q

39. Saravanan R, Holdbrook DA, Petrlova J, Singh S, Berglund NA, Choong YK, et al. Structural basis for endotoxin neutralisation and anti-inflammatory activity of thrombin-derived c-terminal peptides. Nature Communications [Internet]. 2018 Jul;9[1]. Available from: 10.1038/s41467-018-05242-0

40. Miller BR, McGee TD, Swails JM, Homeyer N, Gohlke H, Roitberg AE. MMPBSA.py: An efficient program for end-state free energy calculations. Journal of Chemical Theory and Computation [Internet]. 2012 Aug;8[9]:3314–21. Available from: 10.1021/ct300418h

41. Wan F, Torres MDT, Peng J, Fuente-Nunez C de la. Deep-learning-enabled antibiotic discovery through molecular de-extinction. Nature Biomedical Engineering [Internet]. 2024 Jun;8[7]:854– 71. Available from: 10.1038/s41551-024-01201-x

42. Aziz UBA, Saoud A, Bermudez M, Mieth M, Atef A, Rudolf T, et al. Targeted small molecule inhibitors blocking the cytolytic effects of pneumolysin and homologous toxins. Nature Communications [Internet]. 2024 Apr;15[1]. Available from: 10.1038/s41467-024-47741-3

43. Huang J, MacKerell AD. CHARMM36 all-atom additive protein force field: Validation based on comparison to NMR data. Journal of Computational Chemistry [Internet]. 2013 Jul;34[25]:2135–45. Available from: 10.1002/jcc.23354

44. Abraham MJ, Murtola T, Schulz R, Páll S, Smith JC, Hess B, et al. GROMACS: High performance molecular simulations through multi-level parallelism from laptops to supercomputers. SoftwareX [Internet]. 2015 Sep;1–2:19–25. Available from: 10.1016/j.softx.2015.06.001

45. Hoover WG. Canonical dynamics: Equilibrium phase-space distributions. Physical Review A [Internet]. 1985 Mar;31[3]:1695–7. Available from: 10.1103/PhysRevA.31.1695

46. Nosé S. A unified formulation of the constant temperature molecular dynamics methods. The Journal of Chemical Physics [Internet]. 1984 Jul;81[1]:511–9. Available from: 10.1063/1.447334

47. Parrinello M, Rahman A. Polymorphic transitions in single crystals: A new molecular dynamics method. Journal of Applied Physics [Internet]. 1981 Dec;52[12]:7182–90. Available from: 10.1063/1.328693

48. Essmann U, Perera L, Berkowitz ML, Darden T, Lee H, Pedersen LG. A smooth particle mesh ewald method. The Journal of Chemical Physics [Internet]. 1995 Nov;103[19]:8577–93. Available from: 10.1063/1.470117

49. Hess B, Bekker H, Berendsen HJC, Fraaije JGEM. LINCS: A linear constraint solver for molecular simulations. Journal of Computational Chemistry [Internet]. 1997 Sep;18[12]:1463–72. Available from: 10.1002/(SICI)1096-987X(199709)18:12%3C1463::AID-JCC4%3E3.0.CO;2-H

50. Valdés-Tresanco MS, Valdés-Tresanco ME, Valiente PA, Moreno E. Gmx_MMPBSA: A new tool to perform end-state free energy calculations with GROMACS. Journal of Chemical Theory and Computation [Internet]. 2021 Sep;17[10]:6281–91. Available from: 10.1021/acs.jctc.1c00645

